# An engineered multi-step differentiation program in *Escherichia coli* for self-organized spatial patterning

**DOI:** 10.64898/2025.12.19.695475

**Authors:** Emanuele Boni, Hélène Siboulet, Giacomo Ceracchini, Içvara Aor, Gábor Hollό, Hadiastri Kusumawardhani, Yolanda Schaerli

## Abstract

In nature, complex multicellular structures originate from individual cells containing all essential information for differentiation, patterning and morphogenesis. Synthetic biology enables a bottom-up approach to study these processes by engineering and combining individual modules to progressively increase the system’s complexity. Here, we engineered a multi-step program mimicking cell differentiation in the model prokaryote *Escherichia coli*. Starting from genetically identical cells and without providing any external positional information, we generated autonomous spatial patterns of colonies on a solid surface. We first employed a toggle switch to break population homogeneity (symmetry breaking), stochastically differentiating cells into two subpopulations: senders and receivers. Next, we activated expression of a third reporter in receiver colonies located in close proximity to sender colonies via quorum-sensing based communication (paracrine signaling). Finally, we mimic maturation of the newly emerged population by expressing a fourth reporter via an orthogonal, self-activating, quorum sensing signal (autocrine signaling). The diversity of spatial patterns generated by this multi-step program was accurately captured by simulations of a corresponding mathematical model. Together, these results demonstrate that multi-step differentiation programs can be engineered in unicellular bacteria to drive fully self-organized spatial pattern formation.

## I. INTRODUCTION

Self-organization and self-assembly are basic features of all organisms and are key to the structuring and functioning of macromolecules, cells, individuals, populations, communities and ecosystems [1–3]. A crucial challenge in synthetic biology consists in engineering systems capable of such emergent behaviours, with a twofold objective. On the one hand, in line with Feynman’s ‘build to understand’ principle, recapitulating natural development offers the test bed to challenge hypotheses, prove theoretical principles and shed light on complex phenomena, including developmental diseases [4]. On the other hand, harnessing self-organization and self-assembly offers unprecedented opportunities in the fields of tissue engineering and regenerative medicine [5], distributed cellular computing [6] and engineered living materials [7], underlying the potential of this research area for biomedical, architectural and industrial applications.

Populations of cells, despite sharing identical genetic information, can differentiate and specialize with precision, giving rise to self-organized multicellular organisms [8], or to microbial communities, where distinct phenotypes and properties emerge [9]. Cell fate determination is generally unambiguous despite noisy inputs; moreover, it is stable and self-sustained even upon input removal [4]. The process can be visually represented by Waddington’s landscape of creodes, where a cell starts from an undifferentiated state at the top of an hill and rolls down through forks and bumps, ending in one of several valleys that represent the terminally differentiated states of the initial pluripotent cell [10].

The choice of the differentiation trajectory depends on a variety of input signals and cellular properties that shape the landscape in a unique manner for each cell. They include gene regulatory networks [11–14], environmental cues [15], mechanical forces [16, 17], adhesion [18, 19] and intercellular communication [20] (which can be contact-dependent [21] or based on diffusible signals [22], including morphogen gradients [23]). The emergence of hierarchical structures is driven by the interplay between these different elements [24].

In Waddington’s landscape, a cell goes through several stages of differentiation, ‘losing’ access to potential destinies as the maturation occurs. In fact, the cell-fate determination scheme is sequential (i.e. it requires interdependent steps in succession) and not merely combinatorial (i.e. a combination of events without chronological order) [25]. In particular, the emergence of a subpopulation can produce the signal that triggers further differentiation of the surrounding cells [26, 27].

The emerging picture is that of a flexible, ever-changing, responsive surface, where each cell lineage traces a unique trajectory, that is nevertheless capable of ensuring the robustness and predictability of the development process [28]. For example, stem cells can be cultured under specific conditions that promote self-organization into organoids, three-dimensional miniature versions of organs, that recapitulate the complex structures and functions of real organs [29]. Inspired by this paradigm, researchers wondered whether the ‘cell differentiation landscape’ can be shaped and engineered in order to accurately yet flexibly control the organization and assembly of a synthetic living system.

Sophisticated programs have been engineered in mammalian cells [30, 31], including self-assembled patterns through lateral inhibition thanks to the Delta-Notch system [32], intracellular patterns driven by protein-protein interactions [33], Turing-like patterns generated with a reconstituted Nodal-Lefty pathway [34], and symmetric and polarized 3D structures leveraging contact-dependent signaling and differential expression of adhesive molecules [27]. Remarkably, these systems do not require any external cue, thereby the emerging structures are truly autonomous and self-organized.

While synthetic multi-step differentiation has been widely investigated in mammalian cells, progress in synthetic prokaryotic systems has lagged behind. The reason lies in the ease of repurposing multicellularity elements naturally occurring in mammalian cells [18], compared to the challenge of creating *de novo* tools to engineer multicellularity and cell differentiation in bacteria [19, 22]. Nevertheless, the field has seen notable advances, including various examples of symmetry breaking, the transition of a system from a homogeneous state to multiple distinct states, through different strategies [35–37]. However, synthetic prokaryotic cell differentiation programs predominantly rely on externally added positional information, either in the form of chemical gradients [26], light inputs [38] or in the way cells are spatially arranged [39]. Furthermore, most programs are not truly built upon sequential steps, but are rather limited to 1-step differentiation [26, 36], or they require the researcher’s intervention, for example via the addition of chemical inducers during the cellular program [35].

In the present work, we aimed to address this gap by creating an autonomous multi-step program recapitulating cell differentiation in *Escherichia coli* (*E. coli*) (Figure 1) to generate robust yet unique spatial patterns. By engineering the model prokaryote, we were able to mimic some of the early steps that lead to multicellularity in eukaryotic cells. Briefly, we combined a gene regulatory network for symmetry breaking and cell-to-cell communication for paracrine and autocrine signaling [40]. First, we employed a well-characterized genetic module, the toggle switch (TS) [41], to break the homogeneity within the initial population [11, 37] and stochastically differentiate into two stable subpopulations: green senders and blue receivers. Next, we activated a new molecular program in a subset of receivers that are in close proximity to a sender through intercellular communication [42], implemented via the quorum sensing (QS) system LuxI-LuxR [43]. Finally, the newly emerged population underwent maturation by producing an autocrine signal via the quorum sensing system CinI-CinR [44]. The interconnected differentiation steps created three distinct colony types on an agar surface, resulting in a macro-scale spatial pattern. Throughout the work, experimental results were used to develop a mathematical model, providing us with insights that guided further experimental efforts.

**FIG. 1.**
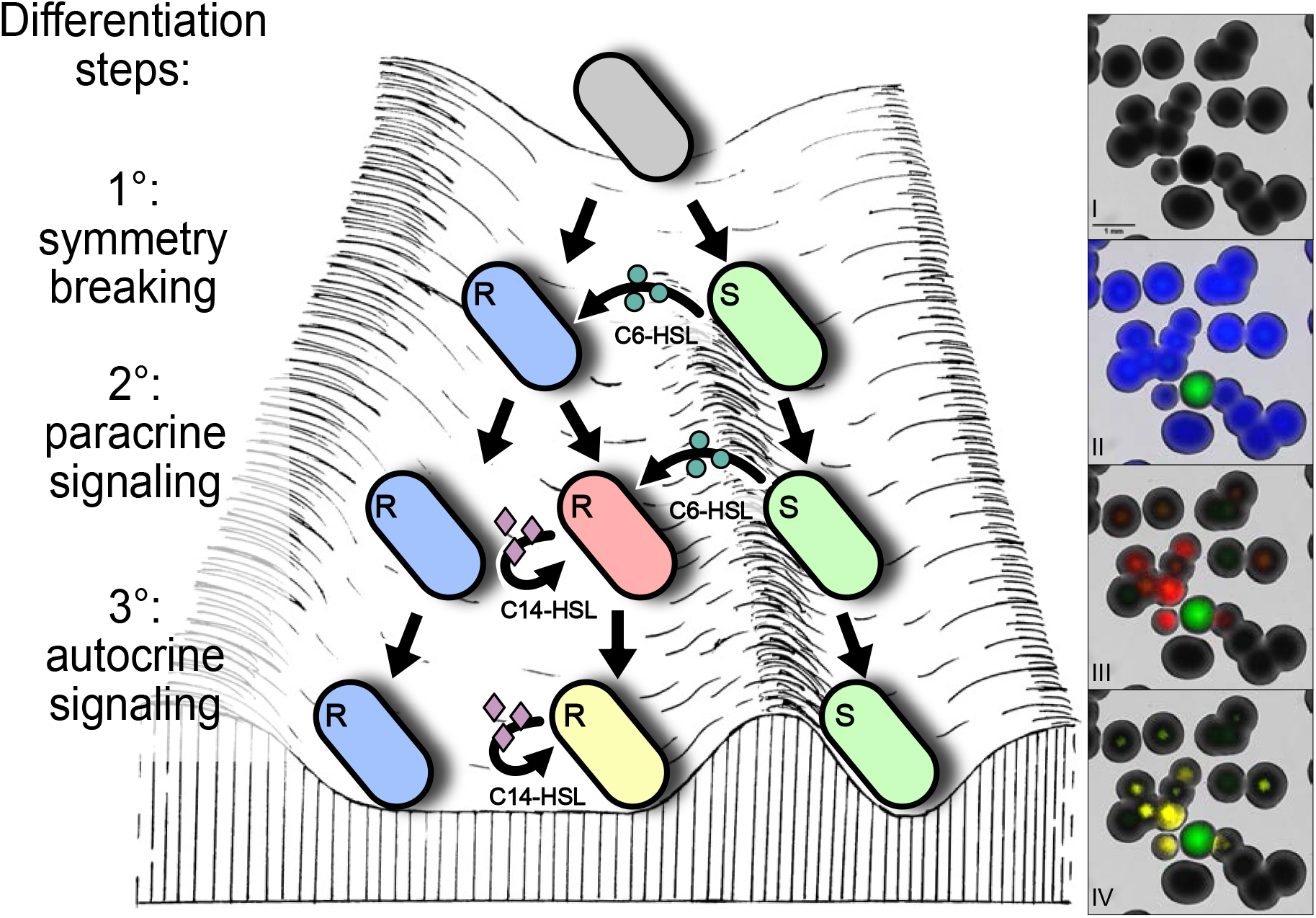
Left panel: graphical representation of a differentiation landscape with three sequential differentiation steps. Green circles: small diffusible molecule 3O-C6-HSL. Purple diamonds: small diffusible molecule 3O-C14-HSL. Cell colour reflects the active molecular program: undifferentiated cells are gray, sender cells are green, receiver cells are blue, receivers exposed to 3O-C6-HSL become red, receivers exposed to 3O-C14-HSL become yellow (while still maintaining their underlying blue identity). Right panel: representative microscopy image of *E. coli* colonies engineered with the 3-step sequential differentiation program, coloured to highlight the three steps. Channels: I - brightfield; II - brightfield, green, blue ; III - brightfield, green, red ; IV - brightfield, green, yellow.

## II. RESULTS

We initially investigated how to differentiate an homogeneous population of *E. coli* cells into two distinct cell types. We leveraged the bistability property of the toggle switch [41], a simple gene regulatory network characterized by two mutually exclusive stable states. We used a TS composed of the transcription factors TetR and LacI, that mutually repress each other by inhibiting the partner’s promoter (pTet and pTrc, respectively). We coupled them to the fluorescent reporters mCerulean and GFP, respectively, by placing the fluorescent proteins under the control of a second pair of pTrc and pTet promoters. Alongside GFP we included LuxI, which will be important later, for the 2*^nd^* differentiation step. TetR and LacI interactions could be tuned by the chemical inducers anhydrotetracycline (aTc) and isopropyl-*β*-D-1-thiogalactopyranoside (IPTG), respectively [45, 46]. IPTG promoted the TetR-mCerulean (blue) state, while aTc drove the system towards a LacI-GFP (green) state (Figure 2a).

**FIG. 2.**
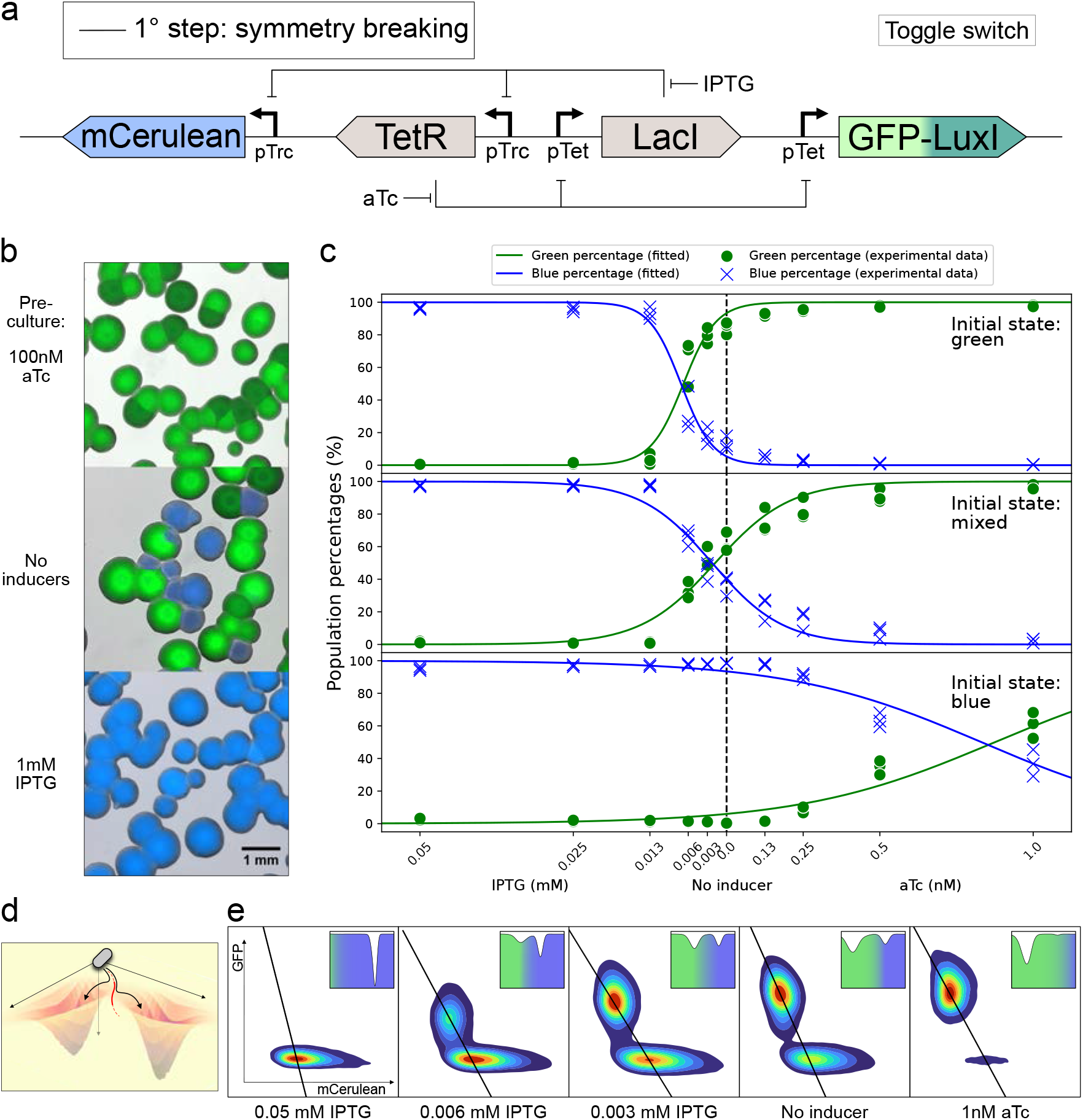
Toggle switch-based 1-step differentiation system. a) Genetic circuit of the 1-step differentiation system, including interactions and chemical inducers (LuxI in the GFP-LuxI fusion protein is not necessary for the 1*^st^* step, but was included, as it will be important for the 2*^nd^* step). Coloured boxes: genes; L-shaped arrows: promoters; T-shaped lines: repressions. b) Microscopy pictures of colonies harbouring the toggle switch. Cells in the initial state mixed were pre-cultured in presence or absence of inducers (indicated on the left side of the picture), then plated and grown overnight in absence of inducers. Channels: bright-field, green, blue. c) Green and blue population percentages in liquid cultures conditioned with different inducer concentrations. Starting from 3 different initial states (green, mixed, blue) cells were pre-cultured in liquid, then the population ratio was quantified via flow cytometry. For the initial state green and blue, cells were taken from colonies that were homogeneously green or blue, while for the initial state mixed, cells were taken from a colony obtained immediately after transformation of the circuit plasmids into cells. Dots represent experimental points (3 biological replicates per condition), lines represent fitted curves (see Methods). Equations and parameters of the fit are indicated in Supplementary Information Table VII. d) Representative 3D energy landscape: valleys represent the green and blue stable states, red line represents the separatrix. The landscape was generated from quantitative flow cytometry data from a single replicate in c (initial state: mixed, induced with 0.003 mM IPTG). We measured GFP and mCerulean fluorescence intensity from 50,000 single cells (see Methods). The depth of each (x,y) point in the landscape corresponds to the number of recorded cells with a specific (GFP, mCerulean) intensity. This specific condition was chosen for illustrative reasons, to show the landscape in an almost symmetrical condition. e) Flow cytometry density plots of 5 representative conditions from c (initial state: mixed, inducer concentration indicated below each plot). For each density plot, we computed the coordinates of the local minima in the green and blue populations, then generated an orthogonal plane crossing them. Black lines represent the projections of these planes on the (GFP, mCerulean) plane. Axes (GFP and mCerulean fluorescence) in the leftmost plot apply to all plots. Inserts represent the 1D energy surface corresponding to each plot, with local minima corresponding to stable states, or valleys. Each 1D plot is the cross section from the corresponding 3D energy landscape (as in d) with the plane crossing the local minima of the green and blue populations, for a single replicate. The colour gradient reflects the inclination of each condition towards the green or the blue state.

With the aim of exploring the self-organization properties of the system, we developed a simple differentiation assay (see Methods, Differentiation assay). Briefly, cells were precultured in liquid medium in presence of different concentrations of chemical inducers, then they were serially diluted and plated on an agar plate without chemical inducers (that is, in absence of any positional information) to form colonies. When cells were pre-cultured with either 100 nM aTc or 1 mM IPTG, the resulting colonies were almost entirely homogeneously green or blue, respectively. When cells were pre-cultured in absence of inducers, both green and blue colonies grew in the plate (Figure 2b) The sporadic appearance of sectors within a colony would pinpoint a stochastic state switch during colony growth (Supplementary Figure S1). In our differentiation assay, cell culturing, dilution and plating correspond to setting the initial conditions for the system. Upon initialisation on the plate, no further intervention was performed, therefore the system evolved in an autonomous fashion.

In order to better characterize the tendency of the TS towards the two stable states, we measured the percentage of green and blue cells in liquid cultures conditioned with a refined range of inducers. We observed three slightly different profiles depending on the initial state of the system (Figure 2c, Supplementary Figure S2), consistently with the TS ‘memory’ property (also called hysteresis [47]). For cells starting from the green or the blue state, the system was biased towards that state, but it could be reversed by adding enough IPTG or aTc, respectively. For cells derived from freshly transformed colonies and pre-cultured without inducer (mixed state), we observed both green and blue cells, with a moderate bias towards the green state (approximately 62:37 green:blue in absence of chemicals). Inducers concentrations as low as 12 *µ*M IPTG and 0.5 nM aTc were sufficient to generate a population close to 100% blue or 100% green, respectively.

Using Waddington’s differentiation landscape framework, we started outlining a multistep differentiation system as a set of valleys and bumps. The TS states correspond to two separate valleys: cells can be in one of the two stable states. In contrast, the saddle (also called ‘separatrix’) between the two valleys represent an unstable undifferentiated state (Figure 2d). In the proximity of the separatrix, the basal transcriptional activity is sufficient to drive the system into a valley (i.e., a stable state) [11]. Mathematically, the profile of the toggle switch landscape corresponds to the number of intersections between the nullclines (the curve where the derivative of a given species over time equals zero), which can be one or three [41]. If they intersect only once, there is only one minimum on the potential curve, which corresponds to a single stable state. If they intersect three times, there are two minima and one maximum, which corresponds to two stable states, and the maximum represents the separatrix. When the system is close to the separatrix, cells can progress towards both the green and the blue destiny. The differentiation trajectory of a single cell is largely stochastic and is influenced by bursts of gene expression. Once the TS is locked in one state, the progeny of that cell maintains a memory of that state, hence a colony has the same state as the founder cell. At the population level, the ratio of green and blue colonies reflects the probability of each cell to fall into the green or the blue valley.

The ratio of green:blue cells at each inducer concentration depended on the position and sharpness of the separatrix, offering a precise mapping of the TS differentiation landscape for each condition (Figure 2e). From a quantitative dynamical-systems perspective, the addition of aTc or IPTG modified the geometric shapes of the nullclines, shifting the three solutions. If the inducer concentration is large enough, the nullclines intersect only once, producing a single stable steady state [41]. Taken together, these observations showed that, owing to the TS bistability, a group of initially undifferentiated cells bifurcated into one of the two possible states, which were stably maintained thanks to the TS hysteresis [48]. We refer to this first differentiation event as ‘symmetry breaking’ (Figure 1).

We subsequently engineered a 2-step differentiation system by combining the toggle switch genetic module with a quorum sensing system. QS systems rely on a synthase (I) for the production of a small diffusible molecule (homoserine-lactone, HSL) and on a receptor (R) for its detection. QS systems differ in the length and chemistry of the HSL acyl chain. Here we used the *Vibrio fisheri* QS system [43], based on 3O-C6-HSL (N-(3-oxo-hexanoyl)-L-homoserine lactone). We coupled each of the two TS states with the production of a QS component: the green state with the synthase LuxI, and the blue state with the receptor LuxR. Cells can therefore exist in two states: green ‘senders’ (S) and blue ‘receivers’ (R) (Figure 3a). Green sender cells, expressing LuxI, produce 3O-C6-HSL, which can diffuse across the cell membrane. Upon entering blue receiver cells, the HSL binds to the LuxR receptor and activates a downstream genetic program, in our case the expression of the red reporter mCherry (Figure 3b). We refer to this second differentiation step, which is subordinate to the first, as ‘paracrine signaling’ (Figure 1), as the diffusible signal influences nearby cells.

**FIG. 3.**
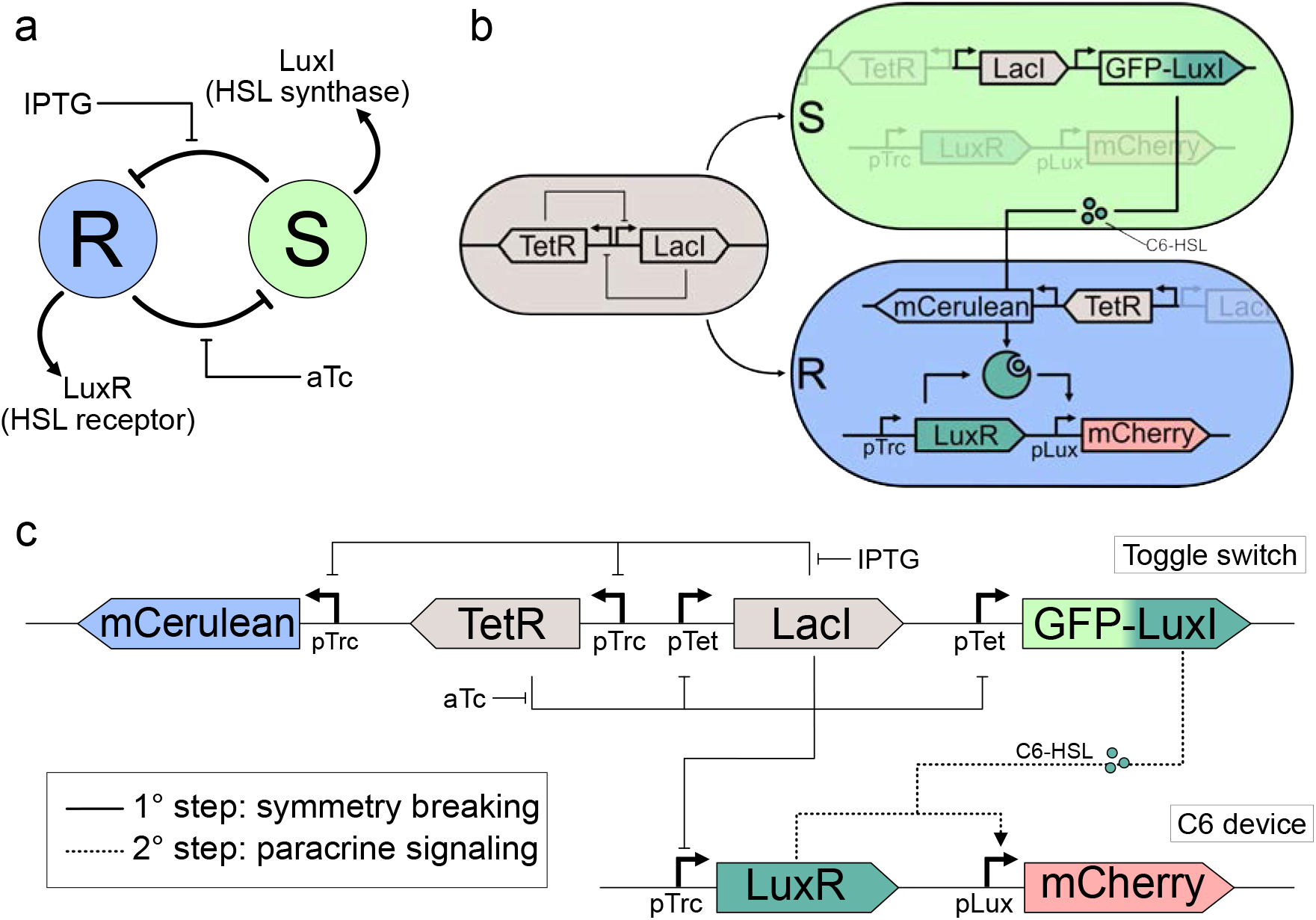
Implementation of a 2-step sequential differentiation system. a) Schematic representation of the 2-step differentiation system. The toggle switch bifurcates the system into two mutually exclusive stable states, receiver (R, blue) and sender (S, green), that are coupled to the two QS components LuxR and LuxI respectively. b) Illustration of the 2-step system functioning: an undifferentiated cell (gray) can become either a green sender or a blue receiver. Upon detection of the 3O-C6-HSL produced by senders, receivers can activate production of the red reporter mCherry. c) Genetic circuitry of the 2-step differentiation system, including interactions and chemical inducers. Coloured boxes: genes; L-shaped arrows: promoters; T-shaped lines: repressions; dotted arrows: activations; green dots: small diffusible molecule 3O-C6-HSL.

It is important to note that all cells in the system contained the same genetic circuitry, composed of two plasmids. In order to include the QS components, we fused the GFP with the LuxI sequence, resulting in a fusion gene. Also, we placed the LuxR gene on a separate plasmid, under control of a pTrc promoter, effectively wiring it to the receiver (TetR-mCerulean) state. On the same plasmid, we placed the red reporter mCherry under control of the pLux promoter (*BBa R*0062 on the iGEM registry), that is targeted by the LuxR:3O-C6-HSL complex. The combination of the receptor pTrc-LuxR and the reporter pLux-mCherry constituted the C6 device (Figure 3c).

In order to characterize the second differentiation step, we first tested reporter cells carrying the C6 device in isolation. Initially, we investigated the profile of mCherry production as a function of 3O-C6-HSL concentration in liquid culture (Supplementary Figure S3). We then moved to solid cultures and observed how colonies harbouring the circuit responded to 3O-C6-HSL (2 *µ*l, 50 *µ*M) spotted in the centre of the plate and diffusing through the LB agar (Figure 4a, top). The red signal was not negligible in absence of 3O-C6-HSL, suggesting that the system presented a degree of leakiness. Using the same assay, we carried out a screening of pLux promoter variants, testing combinations of Lux boxes (G1 [iGem part *BBa K*1216007] and pLux76 [49]) and promoters with different strengths (100% and 54%), in order to identify variants with minimal leakiness and a high fold-change. We identified pLux76 [49] as the regulatory region with the highest fold-change and sufficiently low leakiness among our candidates (Supplementary Figure S4). In the plate assay, the optimized version showed minimal leakiness (Figure 4a, I° row) and, upon induction, a red signal decreasing from the centre to the periphery of the plate, with an improved dynamic range compared to the original pLux version. Colonies growing in the periphery were not red, suggesting that the 3O-C6-HSL level was below threshold and could not trigger mCherry production (Figure 4a, II° row). Subsequently, we characterized 3O-C6-HSL production from green senders. In agar plates, we co-cultured reporter colonies with a sender colony (pre-differentiated with 100 nM aTc and spotted in the centre of the plate). The inducer production from the sender colony gave rise to a pattern of red expression whose intensity was decreasing with the distance from the green sender (Figure 4a, III° row). The induction range obtained with 100 picomoles of pure 3O-C6-HSL was approximately 6-8 mm (50% of maximal induction at 3.74 mm). The induction range around sender colonies was significantly smaller (50% of maximal induction at 1.8 mm, Figure 4b). Mathematical simulations yielded a similar slope when using 0.25 picomoles of 3O-C6-HSL (data not shown), even though care should be used when comparing diffusion of a fixed amount of inducer with continuous production from a growing source.

**FIG. 4.**
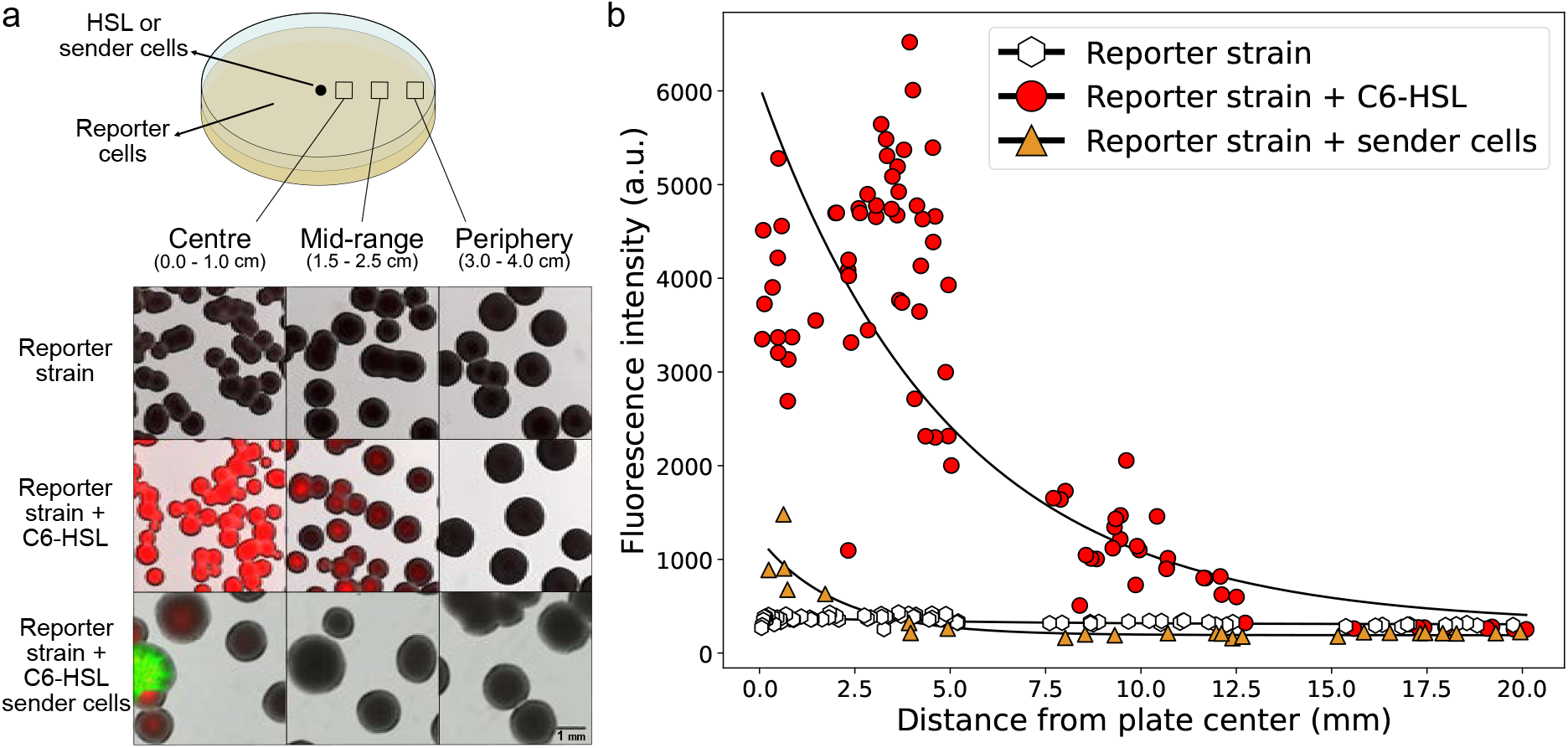
Characterization of the sender and receiver states. a) Top: schematic representation of the plate assay: cells containing only the C6 device were spread homogenously in the plate in order to have well separated reporter colonies. Either pure 3O-C6-HSL or sender cells (pre-cultured with 100 nM aTc) were spotted in the centre of the plate. After incubation, we took images at increasing distance from the centre of the plate. Bottom: microscopy images of reporter colonies growing on plates in absence of 3O-C6-HSL (I° row), in presence of pure 3O-C6-HSL (2 *µ*l, 50 *µ*M, II° row) or of sender cells (III° row, the senders correspond to the green colony). Channels: brightfield, red, green. b) Quantification of red fluorescence intensity from the colonies in a, as a function of the distance from the plate centre. Hexagons, circles and triangles represent the average fluorescence intensity of single colonies; black lines represent exponential curves fitted to experimental points. Equations and parameters of the fit are indicated in Supplementary Information Table VIII.

We then tested the potential of the 2-step differentiation system to generate self-organized spatial patterns. We observed rare motifs arising within the sporadic colonies showing both green and blue sectors. Due to the close proximity of the sender and receiver bacteria within a colony, the blue sectors always showed strong red signal (Supplementary Figure S5). We then generated a well-mixed population of homogenous sender and receiver colonies, by incubating the liquid pre-culture without inducers. Also under this condition, all blue colonies showed high expression of the red reporter, indicating that 3O-C6-HSL reached saturating levels in the plate (Supplementary Figure S6). We therefore selected a condition that would consistently generate a population strongly biased towards the receiver state, with only a few sparse senders to produce the diffusible signal. Having characterized the TS differentiation landscape (Figure 2), we decided to pre-culture cells starting from the green state in presence of 9, 12 and 18 *µ*M IPTG, then plated at a cell density of 500-2000 colonies per plate, in absence of any positional information. This resulted in few sparse green senders densely surrounded by blue receivers, in a highly reproducible manner. The 2-step system autonomously formed a simple pattern composed of green focal points surrounded by blue-red halos, while receivers further away from senders remained blue only (Figure 5a). This motif is reminiscent of a bullseye pattern emerging not within a single colony, but in groups of colonies. Intensity of red expression decreased as the distance from the green sender increased (Figure 5b), while we did not find correlation between the red signal and the colony size (Supplementary Figure S7).

**FIG. 5.**
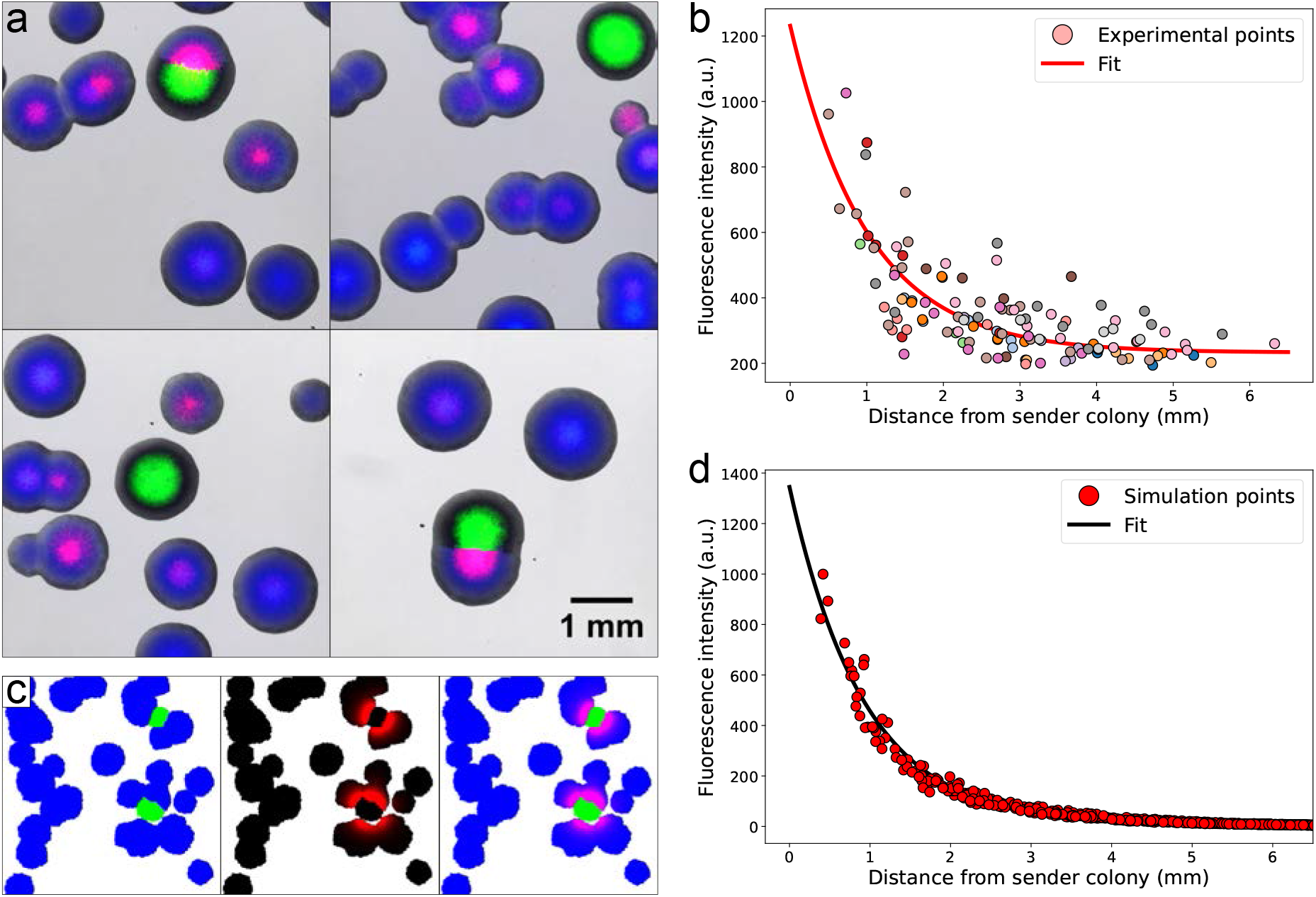
The 2-step sequential differentiation system forms a simple self-organized spatial pattern. a) Microscopy images of the pattern generated by the 2-step system in the immediate surroundings of green sender colonies. Channels: brightfield, red, green, blue. b) Quantification of the red fluorescence intensity of receiver colonies as a function of the distance from the closest sender colony. Each dot represents the average fluorescence intensity of a colony, n = 126 colonies across 14 different images, dots coming from the same image are filled with the same colour. Red line represents an exponential curve fitted to experimental points. c) Mathematical simulation of the spatial pattern at 24 h. Colonies are colour-coded, from left to right, according to: TS state (green-senders, blue-receivers), mCherry fluorescence intensity, full pattern. d) Quantification of the red fluorescence intensity of receiver colonies as a function of the distance from the closest sender colony. Dots come from 10 mathematical simulations; black line represents an exponential curve fitted to simulation points. Equations and parameters of the fit are indicated in Supplementary Information Table IX.

The addition of the QS component increased the differentiation landscape complexity. The receiver valley now widens and contains two regions: one adjacent to the sender valley, and one distant from it. Under the influence of the 3O-C6-HSL produced by the senders, receivers close enough to the senders valley can activate a new genetic program, the expression of the red reporter (in addition to, and not replacing, the blue reporter). Conversely, receivers distant from the senders valley are not exposed to the diffusible signal, so they simply maintain their blue identity (Figure 1, top 3 rows). It is crucial to distinguish that, while the blue/green identity associated with the toggle switch represents a true bistable state [41, 48, 50], the expression of the red reporter is reversible and dependent on proximity to the 3O-C6-HSL source. Although the QS ON-OFF transition is considerably slower than the OFF-ON activation, reversibility remains a fundamental characteristic of QS [51]. Furthermore, protein half-lives in stationary-phase cells extend over tens of hours [52, 53], allowing for the stable detection of mCherry fluorescence over days. Consequently, mCherry serves as a robust readout for the molecular program’s output; however, a truly differentiated cell state would necessitate irreversible modifications to the gene expression profile. Therefore, in our landscape analogy we represented the receiver valley as a continuum where red intensity decreases as the distance from the sender valley increases, without local minima (Figure 1, 3*^rd^* row).

The spatial information provided by the plate assays allowed us to ‘map’ the receiver valley, in relation to the sender valley. Due to the limited amount of diffusible molecule produced by sender cells, HSL-induced red expression only occurred in close proximity (i.e. within 2-3 mm) to the green colonies. Besides the spatial dimension, we were interested in the temporal dynamics of the system, i.e. the speed of the evolution from the undifferentiated state to the terminally differentiated state. We therefore collected time-lapse videos of the plate assay, first inducing receiver cells with pure 3O-C6-HSL, then spotting sender cells alongside receivers, and finally performing the full differentiation assay (Supplementary Movies 1, 2, 3, 4). Whilst one might expect colonies to first appear green or blue and only later acquire red fluorescence, our time-lapse experiments showed that colonies were already expressing the final fluorescent reporters by the time they became visible. Thus, our observations indicate that, at the colony level, HSL production and detection, induction of the red reporter, and colony growth occur concurrently over the same time period. When colonies stopped expanding the fluorescence signal kept increasing (probably due to the maturation of fluorescent molecules expressed earlier), but, despite the fact that 3O-C6-HSL diffusion continued, there was no apparent progression of the red expression front, suggesting that entry in stationary phase marked the terminal differentiation, in agreement with data from liquid cultures (Supplementary Figure S8). When colonies stopped expanding, the fluorescence signal continued to increase (probably due to the maturation of fluorescent molecules expressed earlier), but, despite the fact that 3O-C6-HSL diffusion continued, there was no apparent progression of the red expression front, suggesting that entry into stationary phase marked the terminal differentiation, in agreement with data from liquid cultures (Supplementary Figure S8).

We developed a mathematical model combining spatial and temporal information to qualitatively recapitulate the patterning properties of the system (see Methods, Supplementary Figure S18 and Supplementary Information section A). We started by fitting the experimental results of the TS differentiation space with a sigmoid function (Eq. 2), in order to determine the ratio of green-senders:blue-receivers at any given inducer concentration (Figure 2). Next, we modelled colony growth on solid surface via a cellular automaton (Supplementary Figure S19). We solved the 3-dimensional diffusion equation (SI Eq. 3) numerically to simulate 3O-C6-HSL diffusion in the agar. The mCherry production as a function of the 3O-C6-HSL concentration over time was fitted with a Hill function (SI Eq. 5) using data from liquid culture (Supplementary Figure S3). We accounted for the reduction of bacterial activity over time (i.e. the transition from exponential to stationary phase) by assuming that the biosynthetic capacity of cells is proportional to a monotonically decreasing sigmoid of the time (SI Eq. 7), setting a limit to the production of both fluorescent reporters and HSL synthases (Supplementary Information sections A 3 and A 5). We combined these elements into a coherent spatio-temporal model of the 2-step differentiation system, whose simulations can recapitulate the evolution and final pattern of the system for any given green-sender:blue-receiver ratio (Figure 5c and Supplementary Movie 5) with remarkable qualitative agreement (Figure 5d). Every parameter of the model can be tuned in order to forecast the outcome of experimental variations (e.g. changing the agar thickness or the abundance of colonies; see Supplementary Information section A), including simulating a single sender colony or a droplet of HSL in the centre of the plate (Supplementary Figure S9). A representative simulation of the 2-step differentiation system across multiple time points is shown in Supplementary Figure S22.

We therefore employed the mathematical model to simulate the patterns generated from the 2-step differentiation system for different initial blue:green ratios. The model suggested a variety of outcomes, from 100% green, to 100% blue, passing through intermixed conditions where varying amounts of red colonies appeared, due to the different frequency of senders (hence, to the different amount and distribution of 3O-C6-HSL, Figure 6a). We successfully validated the simulations experimentally, by initiating the system at 11 different initial conditions, corresponding to 11 distinct differentiation landscapes, by changing one single variable: the amount of inducers in the pre-culture (Figure 6b). We observed a remarkable agreement between experimental patterns and simulations with corresponding green:blue ratios (Figure 6c,d). In particular, our experimental data were consistent with the simulations, showing that the entire blue population was also red until the fraction of blue colonies exceeded that of green colonies, after which the red fraction declined to levels comparable to the green population (Figure 6a,b). Due to the intrinsic stochasticity of the system and the random localization of green senders in the plate, each condition gave rise to a predictable yet unique pattern, features that are reminiscent of the robustness and uniqueness underlying development and self-organization in natural systems [54, 55]. Employing the 2-step differentiation system to generate a variety of unique motifs demonstrated the potential of engineering the differentiation landscape to program autonomous spatial patterning in *E. coli*.

**FIG. 6.**
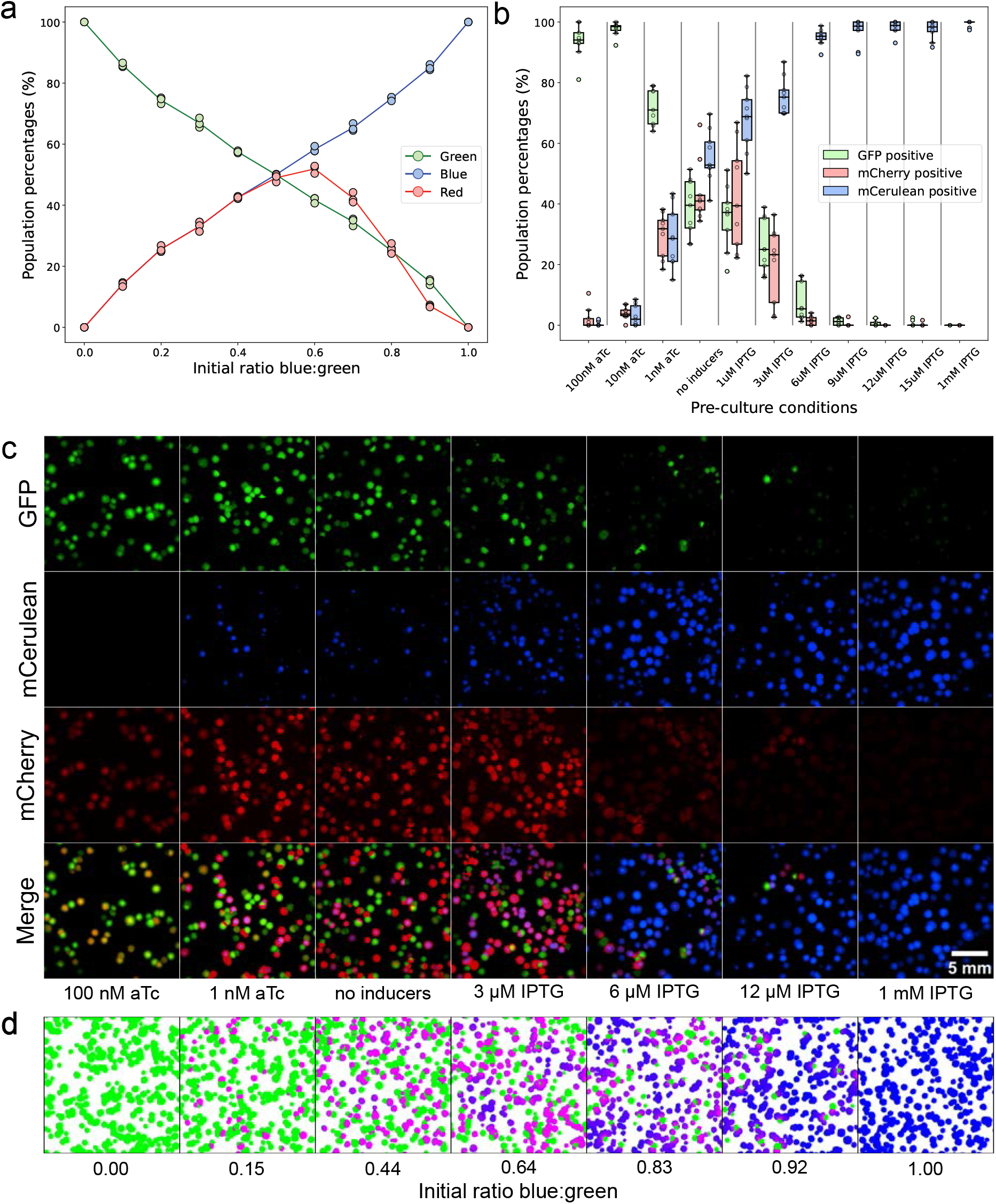
The 2-step sequential differentiation system can generate a variety of spatial patterns. a) Quantification of the three populations abundances from mathematical simulations of the system, in function of the initial blue:green ratio. Dots represent simulation points (n=3 simulations for each initial condition), lines connect averages. b) Quantification of the three populations abundances (green-senders, blue-receivers, red-receivers) in microscopy images. Cells harbouring the 2-step system were pre-cultured with 11 different inducer concentrations (indicated on the x axis). Boxplots of 3 biological replicates (3 images per replicate) represent 1*^st^*, 2*^nd^*, 3*^rd^* quartile of the distribution, whiskers extend to the rest of the distribution or to 1.5 · *IQR* (interquartile range), dots represent individual data points (including outliers). c) Representative microscopy images of the spatial patterns generated by cells harbouring the 2-step system, pre-cultured with different inducer concentrations (indicated at the bottom of each column). Each column displays one representative image (from four locations imaged) of the seven plates in b. Rows (from top to bottom): GFP channel, CFP channel, mCherry channel, composite image. d) Mathematical simulation of spatial patterns at 24 h, starting from different blue:green ratios (indicated below each simulation). Colonies are colour-coded according to their simulated green, blue and red fluorescence intensity. Initial blue:green ratios were chosen to match experimental blue:green ratios in c.

Finally, we extended our program to mimic a maturation step for the newly emerged red population. Thanks to the production of an ‘autocrine signal’ that affects only the red cells, this population drifts apart from the blue receiver state, increasing the distance between the two cell types in the differentiation landscape (Figure 1, IV° row). Similarly to the 2*^nd^* step, the 3*^rd^* step recapitulates a differentiation trajectory via expression of a fluorescent reporter, which in this system is reversible and does not introduce irreversible modifications of the cell state. Concretely, we coupled the expression of the mCherry reporter in blue cells with the production of a second QS synthase producing an orthogonal QS signal. All blue-receivers also expressed the cognate QS receptor. We initially attempted to combine the QS system LuxI-LuxR with LasI-LasR from *Pseudomonas aeruginosa* [56]. Despite previous reports demonstrating a functional combination of the 3O-C6-HSL and the 3O-C12-HSL signals in the context of synthetic circuits [49], our efforts were unsuccessful, primarily due to genetic cross-talk between the LuxR receptor and the pLas81 promoter (Supplementary Figures S10, S11, S12). Therefore, we explored a different QS molecule, with a longer acyl chain: 3O-C14-HSL (N-(3-Oxotetradecanoyl)-L-homoserine lactone), produced and sensed by the regulatory locus cinRI in *Rhizobium leguminosarum* [44]. We observed minimal cross-talk between the two QS systems and therefore combined a C14 device with the C6 device. In the presence of 3O-C6-HSL, the C14 synthase CinI was expressed from the pLux promoter in the same operon containing the reporter mCherry. The receptor CinR was expressed in all blue-receiver cells, and the reporter mCitrine was placed under the control of the pCin promoter (Figure 7a). We expected that different subsets of the full genetic circuitry would be activated in a sequential fashion during the differentiation cascade, namely that mCherry and CinI would be expressed only in blue receivers in close proximity to green senders (Figure 7b). However, we found that leaky expression of CinI from pLux in green sender colonies produced enough 3O-C14-HSL to erroneously trigger mCitrine expression in nearby receivers (Supplementary Figure S13a). Therefore, we replaced the pLux promoter with pLuxLac, which was repressed by LacI in the green-sender state, effectively minimizing the leakage of the promoter and avoiding unwanted production of 3O-C14-HSL (Supplementary Figure S13b). In this optimized circuit, the three steps proceeded in the intended sequence (Figure 7c): blue-receivers in proximity of green senders produced both red and yellow signals (Figure 7d), whose intensity was decreasing as the distance from the closest sender colony increased, with similar slope (Figure 7e).

**FIG. 7.**
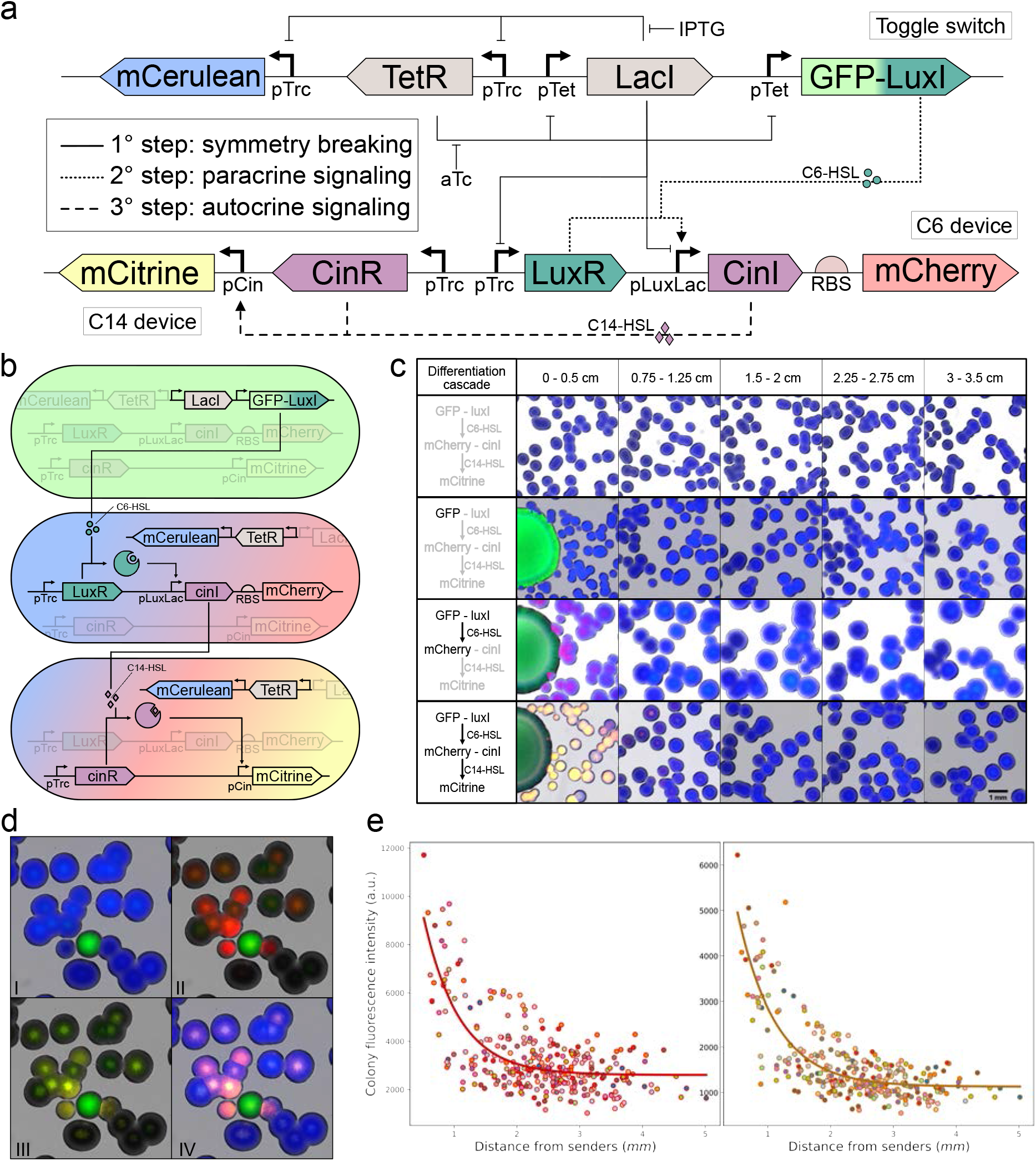
Implementation of a 3-step sequential differentiation system. a) Genetic circuitry of the 3-step differentiation system, including interactions and chemical inducers. Coloured boxes: genes; L-shaped arrows: promoters; T-shaped lines: repressions; dashed and dotted arrows: activations; green dots and purple diamonds: small diffusible molecules 3O-C6-HSL and 3O-C14-HSL, respectively. b) Schematic representation of the 3-step system functioning: a green sender cell produces 3O-C6-HSL via LuxI, which stimulates nearby blue receivers to turn red and to activate production of 3O-C14-HSL via CinI. When the level of 3O-C14-HSL is above threshold, receivers can also turn on production of the yellow reporter mCitrine. c) Demonstration of the 3 sequential steps. The left panel shows the active (black) and inactive (gray) steps of the differentiation cascade. The right panel shows microscopy images of receiver colonies (small, in blue, on the right) grown in presence or absence of sender colonies (big, in green, on the left), with complete or incomplete circuitry, in accordance with the left panel. I° row: receivers only. II° row: senders and receivers, the LuxI synthase is missing from the circuit. III° row: senders and receivers, the CinI synthase is missing from the circuit. IV° row: senders and receivers, complete circuity. Channels: brightfield, red, yellow, green, blue. The sender colonies in this figure do not originate from one single cell, but from 1 *µ*l of a culture of sender cells at OD=1, which explains their larger size (see Methods). d) Representative microscopy image of the 3-step sequential differentiation system. Channels: I - brightfield, green, blue ; II - brightfield, red, green ; III - brightfield, yellow, green ; IV - brightfield, red, yellow, green, blue. e) Quantification of red (left panel) and yellow (right panel) fluorescence intensity as a function of the distance from a green sender colony. Each dot represents the average fluorescence intensity of a colony, n = 304 colonies across 15 different images, dots coming from the same image are filled with the same colour. Red and yellow lines represent exponential curves fitted to experimental points. Equations and parameters of the fit are indicated in Supplementary Information Table X.

Since 3O-C14-HSL can diffuse out from red-blue cells, similarly to 3O-C6-HSL, one might expect mCitrine expression to be triggered in neighbouring blue-receivers cells, even without local CinI and mCherry production. Indeed, bypassing the first and second differentiation steps by co-culturing pre-differentiated red-receivers in presence of C14 reporter colonies confirmed that red colonies produce and release 3O-C14-HSL at nanomolar levels (Supplementary Figure S14). However, since the full cascade lacks any signal amplification, production of 3O-C14-HSL remains confined to cells that also receive the 3O-C6-HSL signal. Mathematical simulations further confirmed that 3O-C14-HSL can function only as an autocrine signal and not as a second paracrine cue, as its diffusion wavefront consistently lagged behind the wavefront of 3O-C6-HSL diffusion (Supplementary Movie 6). Our model suggests that varying the diffusion coefficients of the two diffusible signals could generate more complex patterns, particularly concentric rings where the expression of red and yellow reporters does not overlap (Supplementary Figure S15). Interestingly, the largest region of mCitrine reporter expression is predicted when the diffusion of the first signal is slow, while that of the second is fast. This suggests that sufficient local accumulation of the first signal is required to reach the threshold for production of the second signal, which can then spread further away to produce an outer blue-yellow region. Finally, we also employed our mathematical model to predict the spatial patterns generated by the 3-step system with varying ratios of sender and receiver cells. These predictions generally align with the patterns in Figure 6d, with the addition of yellow patches overlapping with areas characterized by high red density (Supplementary Figure S16).

## III. DISCUSSION

In this project, we have sought to engineer the cellular differentiation landscape as a way to achieve robust, yet unique, autonomous spatial patterning in a population of *E. coli* cells (Figure 1). While most previous studies have focused on bacterial lawns [13, 23, 49, 50, 57] or on pattern formation within individual colonies [12, 14, 35, 58–60] and cell aggregates [19], here we explored spatial pattern formation across physically separate colonies, adding a distinct scale to the field of synthetic patterning. We first started from a simple 2-valley system, represented by the toggle switch (Figure 2). Then, we remodelled the blue-receiver valley, so that cells in close proximity to the green-sender valley activated a new cellular program (Figures 3, 4). We explored the autonomous patterning properties of the 2-step sequential differentiation system (Figures 5, 6). Finally, we engineered the newly emerged population to diverge further from the blue population, mimicking maturation through autocrine signaling (Figure 7). We also developed a mathematical model to recapitulate the spatial arrangement and identity of colonies on an agar plate and to gain insights into the features of the system (Figures 5b,d and 6a,d).

Three elements contribute to the uniqueness of each cell differentiation trajectory in our engineered system. First, the inducer concentration in the liquid culture translates into different landscapes. Second, the previous state of the system (green or blue or mixed) corresponds to a specific location in the landscape, which can be very close to the separatrix or strongly biased towards one of the two TS states. Third, the stochasticity of the system (i.e., the identity and the spatial arrangement of colonies on the plates) implies that although *n* experiments with identical initial conditions will result in predictable ratios of the different cell types, they will nonetheless produce *n* unique spatial patterns. This effectively recapitulates some of the underlying properties of cell differentiation and self-organization in natural organisms [54, 55]. By using *E. coli* bacteria, that are quintessentially unicellular, to differentiate an homogeneous populations of cells into three distinct cell types, we have strengthened the evidence that synthetic biology is a powerful and versatile platform to engineer complex differentiation programs and explore multicellularity and morphogenesis [4].

Our project advances the field of prokaryotic synthetic differentiation, which has always lagged behind developments in eukaryotic systems. Impressive progress has been made in engineering cell differentiation and pattern formation in bacteria [24, 61] using distinct mechanisms including positional information [26], asymmetric plasmid partitioning [35, 36], recombinase-mediated irreversible cell differentiation [62], quorum sensing [63, 64], motility-based patterns [58, 59] and reaction-diffusion patterns [60]. Yet, few exploited key principles of autonomous multistep sequential cell differentiation to generate self-organized structures. In this work, we successfully engineered a multistep program mimicking the differentiation of an initially homogeneous population into three distinct cell types without any external cues, while still achieving fine-tuning of the populations’ ratios through pre-culture conditions.

Our 3-step sequential differentiation program leverages three key developmental biology mechanisms: symmetry breaking, paracrine signaling and autocrine signaling. In future work, the system could be modified in many ways, depending on the desired pattern and its features. For example, the autocrine signal could be converted into a second paracrine signal, by ensuring it diffuses further than the first. Active degradation of the first HSL via the quorum quenching lactonase AiiA [65, 66], a positive feedback loop that exponentially enhances 3O-C14-HSL production, or choosing signals with significantly different diffusion rates could enable the diffusion wavefront of the second molecule to overtake the first. Natural cell differentiation is typically irreversible and remains stable even after the removal of the trigger [4]. In our system the pattern becomes fixed upon entry in stationary phase, and the toggle switch provides memory for the green and blue states. However, if red cells are re-streaked, they lose their identity unless they remain in proximity to sender cells. The implementation of positive feedbacks or stable genetic modifications (e.g., excision of a DNA cassette [62, 67]) could make our program also irreversible.

Beyond expressing fluorescent proteins, differentially targeting genes involved in cell morphology and growth could drive morphogenetic changes in one (or more) subpopulation(s). Coupling the detection of QS molecules to cell motility [58] could enable fascinating forms of hierarchical patterning. Another promising direction would be to assign distinct functions to each differentiated cell type [35]. Division of labour is a central feature of natural microbial communities and is increasingly being exploited in synthetic biology for a broad range of applications [68]. Spatial gradients, cell–cell communication, and engineered pattern formation have, for example, been harnessed for biological computation and logic processing [69–71]. Engineered microbial communities have also been developed as biosensors [72, 73], and the ability to store information in the form of spatial patterns with defined population ratios could enable environmental biosensors that record past chemical exposures [74]. Finally, in the emerging field of engineered living materials, patterning strategies are being explored to generate materials with self-organized functional architectures [7]. By establishing design principles for synthetic differentiation and spatial self-organization, our work contributes to enabling such functional specialization in engineered systems.

## IV. METHODS

### Media

Lysogeny broth medium (LB: 10 g Bacto-Tryptone, 5 g yeast extract, 10 g NaCl per 1 l) and LB agar plates (agarose 1.5% w/v) supplemented with the appropriate antibiotics (50 *µ*g/ml kanamycin and/or 50 *µ*g/ml spectinomycin) were used for cloning and experiments. For microplate reader experiments, EZ medium (Teknova) containing 0.5% glycerol was used.

### Reagents

Oligonucleotides and chemicals were purchased from Sigma-Aldrich. Polymerase chain reactions (PCRs) for cloning were performed with Phanta Max Master Mix (Dye Plus) DNA Polymerase (Vazyme). Colony PCRs were performed with Taq Master Mix DNA Polymerase (Vazyme). PCR products were purified with the Monarch PCR & DNA Cleanup Kit (NEB). Plasmids were purified using the QIAprep Spin Miniprep Kit (QIAGEN). Gibson assemblies were performed with NEBuilder HiFi DNA Assembly Master Mix (NEB).

### Cloning

Molecular parts were assembled into plasmids via Gibson Assembly cloning. The toggle switch motif was cloned from an already functional toggle switch plasmid (pKDL071) [48], kindly provided by Jeong Wook Lee. The sequences of the receptor *luxR*, the promoters *pLux* and *pLuxLac* [50], the reporter-synthase fusion protein *GFP-luxI*, as well as the fluorescent reporters *mCerulean*, *mCherry* and *mCitrine* [13], were taken from our’s group previous work. The sequences of the synthase *lasI* and the receptor *lasR* were taken from the iGEM registry (*BBa C*0078 and *BBa C*0079, respectively), codon-optimized for *E. coli* with the IDT codon optimization tool and synthesized from IDT. The sequences of the synthase *cinI* and the receptor *cinR* were kind gifts from Matthew Bennett (Addgene plasmid #141124 (C332) [75]) and from Christopher Voigt (Addgene plasmid #108535 [76]), respectively. All plasmids and primers used in this work are listed in Supplementary Information Tables IV and VI, respectively.

### Strain and growth conditions

Plasmids were transformed into *E. coli* str. K-12 substr. MG1655 (NCBI:txid511145). Single colonies were used to inoculate LB medium, grown overnight at 37 °C and 200 rpm shaking in culture tubes, to prepare stocks. Strains were then streaked on LB agar plates and incubated overnight at 37 °C, then plates were stored at 4 °C up to 4 weeks.

### Differentiation assay

Where appropriate (i.e., when it was important to know whether the previous state of the TS was green or blue) the identity of colonies was assessed with a LED based transilluminator (Nippon Genetics). Single colonies were picked from LB agar plates, resuspended in 100 *µ*l sterile PBS, then 10 *µ*l were used to inoculate 1 ml of LB supplemented with antibiotics and the appropriate inducers in 1.5 ml Eppendorf tubes. Unless otherwise stated, a concentration of 1 mM IPTG was used to shift the entirety of the population towards the blue-receiver state, 100nM aTc for the green-sender state, and 9-18 *µ*M IPTG, starting from initial state green, to generate a mixed population with abundant receivers and few senders. Cultures were grown at 37 °C and 200 rpm shaking for approximately 6 h, then their OD was measured with a NanoDrop One (ThermoFisher). Cultures were diluted to OD=1 with sterile PBS, then serially diluted 3 times with sterile PBS, until OD=10*^−^*^3^. Unless otherwise stated, 10 *µ*l of the OD=10*^−^*^3^ dilution were plated and spread out with glass beads on LB agar plates supplemented with antibiotics and no inducers. Where appropriate, HSL or sender cells were spotted in the centre of the plate, and kept on the bench until liquid absorption. Unless otherwise stated, 2 *µ*l of HSL 0.05 mM and 1 *µ*l of sender cells from a liquid culture at OD=1 were used for induction on plates. Plates were incubated upside-down at 37 °C for up to 24 h, then imaged.

### Flow Cytometry

Single colonies were picked from LB agar plates, resuspended in 100 *µ*l sterile PBS, then 2 *µ*l were used to inoculate 150 *µ*l of LB supplemented with antibiotics and the appropriate serially diluted inducers in a 96-well plate. Cultures were grown at 37 °C and 200 rpm shaking for approximately 6 h, then analysed with a CytoFLEX S flow cytometer. We used 488 nm excitation laser in combination with FITC filter (emission 525/40 nm) for GFP measurements and 405 nm excitation laser in combination with PB450 filter (emission 450/45 nm) for mCerulean measurements. Flow cytometry raw data were recorded using CytExpert software (version 2.4). First, to discriminate between cells and other particles, all measured events were gated by forward scatter height (FSC.H) *>* 10^3^ arbitrary unit (a.u.) and side scatter height (SSC.H) *>* 10^3^ a.u. Second, we excluded doublets by plotting the FSC.H against the forward scatter area (FSC.A) and set a gate for the events with approximately 1:1 ratio of FSC.H to FSC.A. We recorded 50,000 events of singlet cells. Data was processed with FlowJo (version 10.8.1). To quantify the percentages of GFP-positive and mCerulean-positive cells shown in Figure 2c, we set gates below and above the y=x diagonal of the green (FITC-H) versus blue (PB450-H) graph, ensuring a signal of at least 10^2^ a.u. in the corresponding channel and at most 10^3^ a.u. in the opposite channel. For a visual representation of our gating procedure, see Supplementary Figure S17. Python (version 3.8.12) was used for data analysis and visualization. Energy landscapes were constructed using Gaussian kernel density estimation (KDE) of the flow cytometry data. Specifically: a) the inverse of the KDE was plotted to generate the 3D landscape (Figure 2d); b) the KDE values were represented as heatmaps to produce 2D landscapes (Figure 2e); and c) a plane intersecting the lowest minima of the regions below and above the *y* = *x* diagonal (bottom-right and top-left portions of the 2D landscape, respectively) was identified, and the KDE cross-section along this plane was plotted to obtain the 1D landscapes (Figure 2e, insets).

### Microplate reader experiments

For Supplementary Figures S3, S8, S10b,c, S11b, S12b and S21, single colonies were picked from LB agar plates, resuspended in 50 *µ*l sterile PBS, then 1 *µ*l was used to inoculate 200 *µ*l of EZ medium (Teknova) containing 0.5% glycerol, supplemented with antibiotics and the appropriate inducers, in a Microtest Plate 96 Well (Sarstedt). The plate was covered with the supplied lid and were incubated at 37 °C with double-orbital shaking in a Synergy H1 microplate reader (Biotek) running Gen5 v3.11 software. OD (absorbance at 600 nm) and fluorescence were measured every 10 minutes for 48 h. Fluorescence settings: mCherry (Ex. 579 Em. 616 - gain 100), mCitrine (Ex. 500 - Em. 541 - gain 65), GFP (Ex. 479 - Em. 520 - gain 65), mCerulean (Ex. 430 - Em. 491 - gain 65). Unless otherwise stated, three biological replicates were measured for each sample. OD and fluorescence values from a blank sample were subtracted from the OD and fluorescence signal of each sample and, where appropriate, the fluorescence value of each sample was normalized by its absorbance at 600 nm to account for differences in bacterial concentration. Python (version 3.8.12) was used for data analysis and plotting.

### Image collection

Microscopy close-up images were collected with a Zeiss Axio vertical microscope running VisiView software, with a 2.5x objective using a brightfield channel and DsRed, YFP, GFP and CFP fluorescent filters. Unless otherwise stated, exposure times were 50 ms for brightfield and 100 ms for the fluorescence channels. When collecting microscopy images with senders and receivers, we carefully avoided to image fields of view with green senders just outside the edges of the images. Where appropriate, 5 images of the same plate, equally spaced by approximately 7.5 mm, were collected along a radius of the plate, from the centre to the periphery, and were subsequently stitched with the appropriate order and orientation. Microscopy wide-field images were collected with a Nikon SMZ25 stereo microscope, with a 1x objective and a 0.63x zoom, using mCherry, GFP and DAPI fluorescent filters. Exposure times were 100 ms for GFP, 200 ms for mCherry, 5 s for mCerulean (DAPI). A dark background was used to minimize diffused light. Microscopy time-lapses were collected with a Zeiss LSM 900 Confocal microscope equipped with incubation chamber and humidity control, running Zen Blue software, using a Plan-Apochromat 10x/0.45 Ph 1 M27 Air objective. Plates were imaged using light (transmitted light, ESID photodiode, 400 nm), red (ex 561 nm, em 560-700 nm), green (ex 488 nm, em 410-546 nm) and blue (ex 405 nm, em 410-546 nm) channels, with a 46 *µ*m pinhole. For the time-lapses, approximately 35 ml of LB agar were poured into the lid of a rectangular plate, then let to dry in a flow hood until use. For plating, 10 *µ*l of cells at OD=10*^−^*^3^ were spotted on the plate, 200 *µ*l of sterile PBS were added, and the droplet was spread homogeneously all over the plate with a T-loop. The plate was kept on the bench until liquid absorption and, where appropriate, HSL or sender cells were spotted in the centre of the plate (either immediately before incubation, or 8 h after incubation started). A small hole was punched on the surface of the agar, next to the plate centre, to help adjusting the focus. Plates were incubated with temperature and humidity control (37 °C and RH 95% respectively) and imaged every 15 min for 48 h, over 20 (10×2, Supplementary Movies 1,2,3) or 81 (9×9, Supplementary Movie 4) adjacent tiles and 3 z-stacks (at 350 *µ*m distance). Tiles stitching was automatically carried out using the Zen Blue software. Despite the humidity control, we observed a shrinking effect in the agar plates, whose thickness passed from ∼3 mm to ∼2 mm.

### Image analysis

Microscopy images were post-processed and analysed with Fiji ImageJ software (version 2.16.0). A custom-made macro was developed to segment colonies based on the brightfield image (combining watershed separation and manual correction), then extract the mean fluorescence intensity of the fluorescent channels, alongside the position of the colony centroid. The latter was used to compute the distance of each colony from the closest green-sender colony or from the plate centre. Calculation of the distance between receiver and sender colonies was performed only for images collected from plates with few sparse (i.e., more than 1 cm away) senders, ensuring that each receiver only sensed the 3O-C6-HSL produced by a single sender colony. Due to bleed-through of GFP signal in the mCitrine channel in the Zeiss Axio microscope, mCitrine signal of green senders was manually set to zero, to overwrite non-existent yellow signal (Figures 1, 7c,d). Python (version 3.8.12) was used for data analysis and plotting. Patterns from mathematical simulations were analysed with Python, applying the same principles. Colonies in the blue and green channel were segmented (using watershed separation) separately, then the mean red intensity was extracted, alongside the position of the colony centroid. The latter was used to compute the distance of each colony from the closest green-sender colony or from the simulation centre. Both image analysis scripts are available on GitHub, with appropriate description of their functioning and requirements, at: https://github.com/SchaerliLab/Multi-step-differentiation-program.

### IPTG–aTc equivalence

IPTG biases the toggle switch towards the blue state, whereas aTc drives it towards the green state. We sought to determine whether IPTG could be mapped onto an effective aTc concentration, enabling us to describe the transition between blue and green with a single continuous control variable. As shown in Figure 2c, a simple linear transformation between IPTG and aTc achieves this for our experiments. The effective aTc concentration ([aTc]_eff_) associated with an IPTG level ([IPTG]) is given by

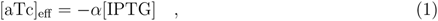

where *α* is a linear conversion factor. The corresponding green fraction *G* was then computed using a sigmoidal response function:

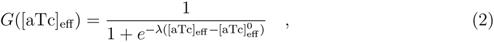

where *λ* denotes the response steepness and [aTc]^0^ the effective midpoint concentration. The blue fraction follows directly from conservation, *B* = 1 − *G*. A single global conversion parameter, *α* = 20 nM/mM, provided an excellent fit to the experimental data (Figure 2c).

**TABLE I.** Fitted parameters (*λ*, [aTc]^0^_eff_ (nM)) describing the toggle-switch response in Figure 2c under different initial induction states (blue, mixed, and green). The linear IPTG–aTc conversion in Eq. 1 used a common parameter of *α* = 20 nM/mM.

| Induction | $\lambda$ | $[\text{aTc}]_{\text{eff}}^0$ |
| --- | --- | --- |
| Blue | 0.1969 | 13.9961 |
| Mixed | 0.5142 | -0.5937 |
| Green | 1.1714 | -2.2507 |

### Mathematical modelling and simulations

Figure S18 summarizes the workflow used to parametrize and construct the model describing spatio-temporal pattern formation. Colony expansion was captured using a cellular automaton framework (SI Section A 1), while intercellular communication via small diffusible molecules was modelled using Fick’s second law (SI Section A 2). Blue receiver cells activated mCherry expression upon sensing 3O-C6-HSL; the mCherry production rate was quantified using plate-reader assays (SI Section A 3). Bacterial activity and viability decreased over time; these effects were incorporated through an experimentally derived activity function, obtained from time-lapse experiments (SI Section A 4). C6-HSL was produced by sender cells; we extracted the C6-HSL production rate from time-lapse microscopy by using the previously calibrated mCherry response curve as a reporter of the C6-HSL concentration (SI Section A 5). In the presence of 3O-C14-HSL, blue receiver cells expressed mCitrine; its production dynamics were characterized via plate-reader measurements (SI Section A 6). The transition towards the yellow state occurred when red cells produced C14-HSL; we estimated the C14-HSL production rate (SI Section A 7), completing the parametrization required for the full spatial model.

## AI Usage Disclosure

During the preparation of this work the authors used AI tools (DeepL, ChatGPT) for language editing. After using these tools, the authors reviewed and edited the content as needed and take full responsibility for the content of the published article.

## AUTHOR CONTRIBUTIONS

E.B. G.C., I.B. and Y.S. designed the experimental research. E.B. and G.C. performed the experiments. I.B, H.K. and Y.S. provided guidance and supervision for the experimental research. E.B. analysed the data and prepared the corresponding figures. H.S. performed the mathematical modelling and prepared the corresponding figures. G.H. provided guidance and supervision for the mathematical modelling. E.B., H.S., G.H. and Y.S. wrote the manuscript. All authors have given approval to the final version of the manuscript.

## DATA AND CODE AVAILABILITY

The plasmids used to generate the main figures in this study and their annotated sequences are available through Addgene at: https://www.addgene.org/Yolanda_Schaerli/ (IDs 251146 to 251150 , see Supplementary Table IV). The source data for all graphs is provided as Extended Data. Raw microscopy images and time-lapses are available on Zenodo: DOI 10.5281/zenodo.17802362. The computer code used for the mathematical simulations of the system and the image analysis scripts used in this study are available on GitHub at: https://github.com/SchaerliLab/Multi-step-differentiation-program.

## Supporting information

Description supplementary movies

## ACKNOWLEDGEMENTS

This work was funded by the Swiss National Science Foundation (grant 310030 200532 awarded to Y.S), and the University of Lausanne.

