## Supplementary information for "An engineered multi-step differentiation program in *Escherichia coli* for self-organized spatial patterning"

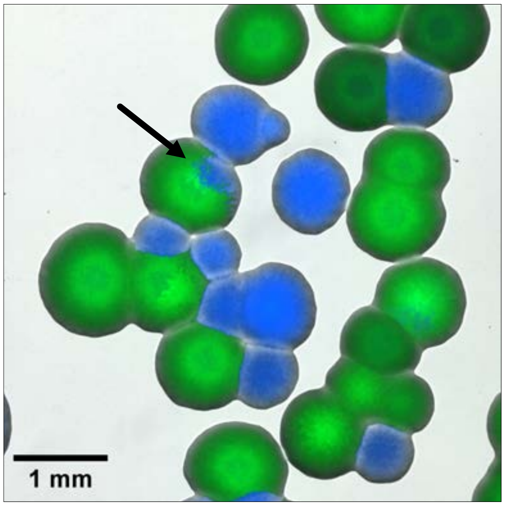


Fig. S1. **Toggle Switch state transitions during colony growth are sporadic.** Microscopy image showing homogeneous and mixed colonies. The Toggle Switch circuit was transformed in *E. coli* cells, a resulting colony was resuspended and grown in absence of inducers, then plated on solid surface in absence of inducers, incubated for 24 h, then imaged. Most colonies were homogeneously green or blue, indicating that all daughter cells inherited the same state as the founder cell. In few cases, colonies with sectors of both colours (marked with a black arrow) could be observed, indicating that a state switch had happened during colony growth. Channels: brightfield, green, blue.

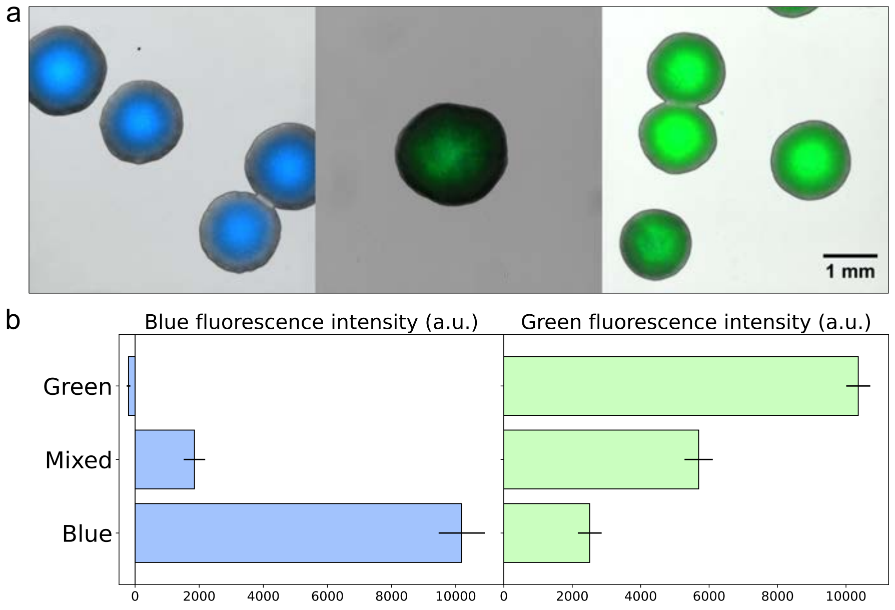


Fig. S2. **Comparison of cell initial states.** a) Microscopy images of colonies containing the toggle switch circuit, pre-differentiated with 1 mM IPTG in the blue state (left), plated directly upon transformation (centre) and pre-differentiated with 100 nM aTc in the green state (right). Channels: brightfield, green, blue. b) Quantification of blue and green fluorescence intensity from colonies in the state green (n=86), mixed (n=54) and blue (n=32), after subtraction of a non-fluorescent negative control (n=174). For each state, colonies from at least 3 different plates were used for the quantification. Bars represent the mean and standard error.

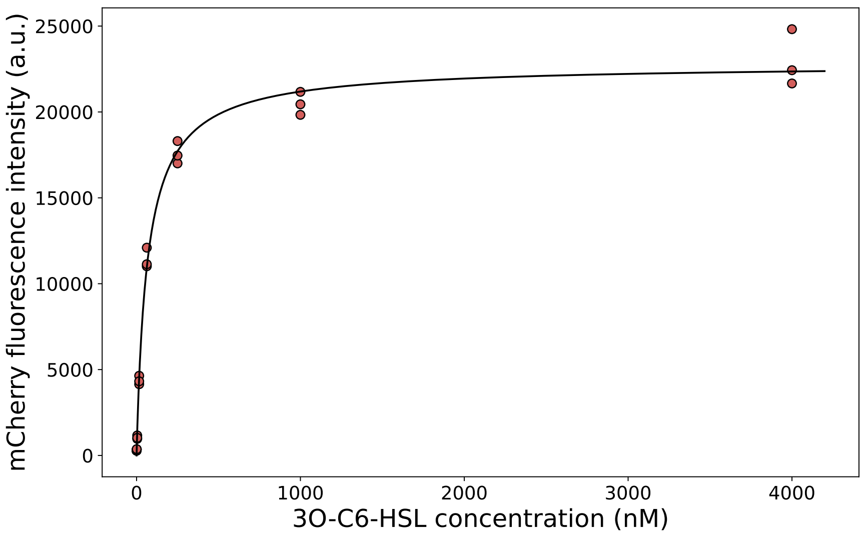


Fig. S3. **Characterization of mCherry fluorescence intensity as a function of 3O-C6-HSL concentration.** Cells containing only the C6 device (LuxR_pLux-mCherry) were grown in a Biotek plate reader for 48 h in medium conditioned with different concentrations of 3O-C6-HSL. Points represent the end-point red fluorescence intensity of 3 biological replicates per condition; black line represents a Hill function (Eq. 5) fitted to experimental points (Equation: $y = \frac{a}{b+x^c}$. Parameters: $a$ = 407.294, $b$ = 0.018, $c$ = -0.955. Method: Levenberg-Marquardt algorithm).

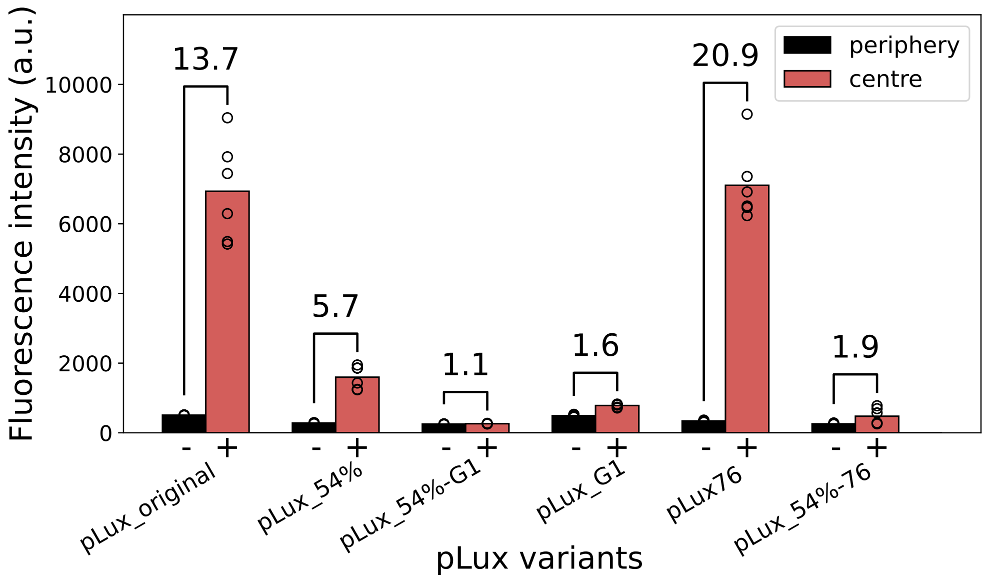


Fig. S4. **Screening of pLux variants.** A small library of pLux variants was created by combining promoters with different strengths and luxO binding boxes with different affinities for the receptor LuxR. The library was tested in solid culture, by plating cells containing the C6 device (LuxR_pLux-mCherry) homogeneously in the plate, then spotting 2 $\mu$l 3O-C6-HSL 50 $\mu$M in the centre of the plate. The promoter efficiency was quantified as the dynamic range (marked on top of each variant) between induction level in the periphery (3.0-4.0 cm from plate centre, bars marked with '-') and induction level in the centre (0.0-1.0 cm from plate centre, bars marked with '+'). Bars represent averages, empty dots represent experimental points (n = 6 colonies per condition). The pLux76 variant was identified as the best candidate from this screening.

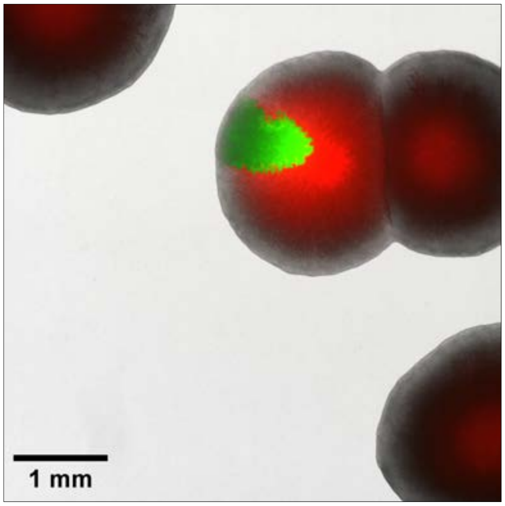


Fig. S5. **The 2-step differentiation system gives rise to intra-colony patterns.** When a colony presented both green and blue sectors, the blue sector was also entirely red. Channels: brightfield, green, red.

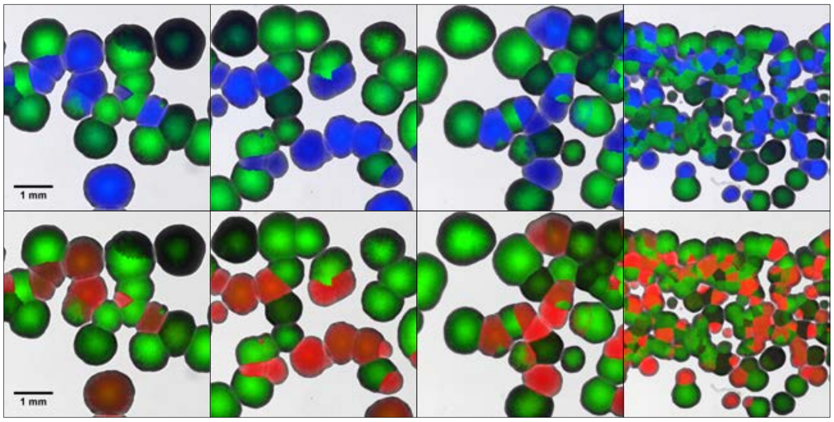


Fig. S6. **Microscopy images of the 2-step sequential differentiation system pattern arising from high intermixing of green-sender and blue-receiver colonies.** Channels: top row - brightfield, green, blue; bottom row - brightfield, green, red. Cells in the initial state 'mixed' were pre-cultured in absence of inducers, then plated on solid surface in absence of inducers, resulting in high intermixing of green and blue colonies. All blue colonies also showed high red signal, indicating an high level of 3O-C6-HSL.

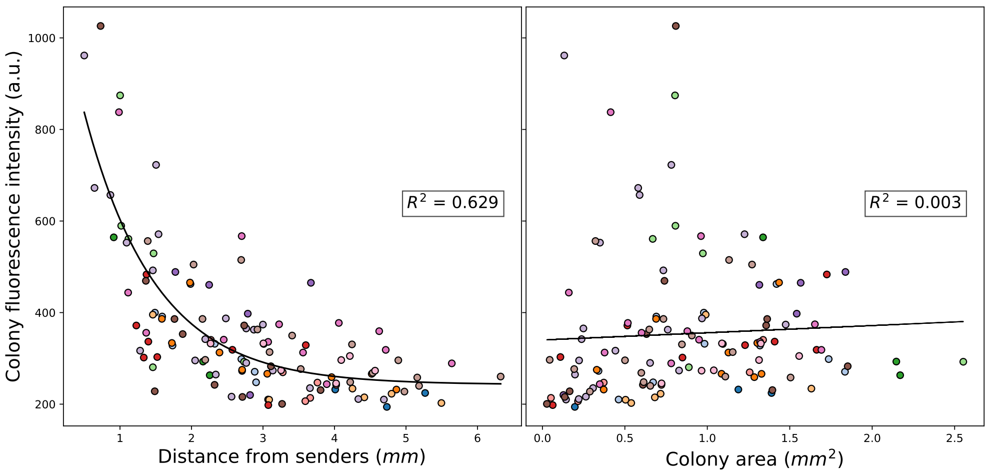


Fig. S7. **Red fluorescence intensity in receiver colonies correlates with distance from sender colonies, but does not correlate with colony size.** Red fluorescence intensity of each receiver colony was plotted against the distance from the closest sender colony (left) and against its own colony size (right). Each dot represents the average fluorescence intensity of a colony, n = 126 colonies across 14 different images, dots coming from the same image are filled with the same colour. Black lines represent an exponential curve (left, equation: $y = \frac{a}{b+x^c}$, parameters: $a = 1.00$, $b = 1223.35$, $c = 242.664$, method: Trust Region Reflective algorithm) and a linear regression (right, equation: $y = a \cdot x + b$, parameters: $a = 15.748$, $b = 340.204$) fitted to experimental points.

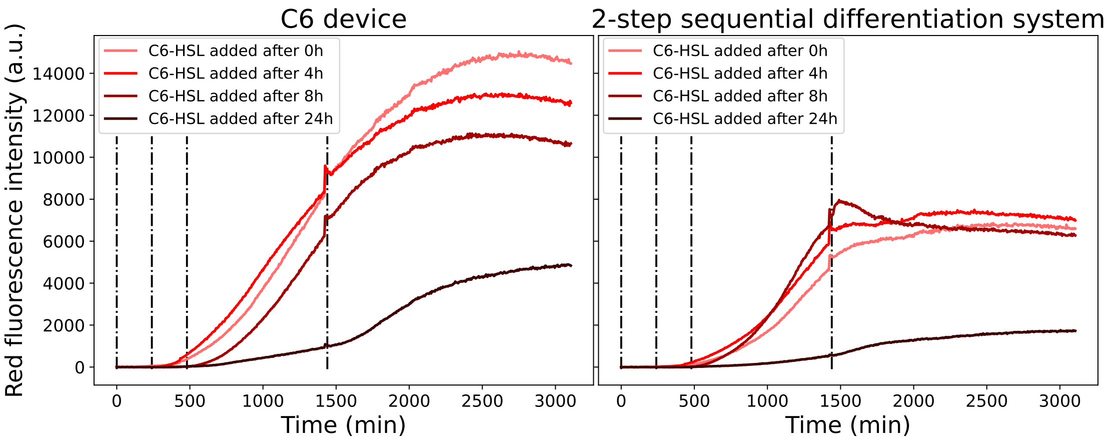


Fig. S8. **Effect of cell growth phase on responsiveness to 3O-C6-HSL.** Cells harbouring the C6 device (left) and the full 2-step sequential differentiation circuitry (right) were grown in a Biotek plate reader with shaking for 48 h and induced with 5 $\mu$M 3O-C6-HSL after 0 h, 4 h, 8 h and 24 h. Early (0 h) induction seems to impose a moderate burden to the cells, especially in the full system. Conversely, late (24 h) induction results in little to no response, probably owing to the entry of cells in stationary phase.

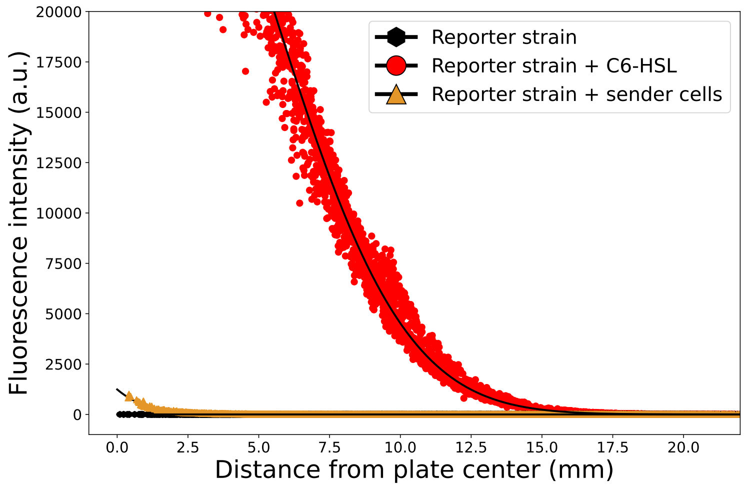


Fig. S9. **Mathematical simulations of the 2-step differentiation system in presence and absence of 3O-C6-HSL.** Quantification of red fluorescence intensity from simulated colonies harbouring the 2-step differentiation system in absence of 3O-C6-HSL (black hexagons), in presence of 50 $\mu$M 3O-C6-HSL (red dots) and in presence of a sender colony (orange triangles), reproducing the conditions in Figure 4 (n=10 simulations per condition). Hexagons, circles and triangles represent simulation points; black lines represent exponential curves fitted to simulation points. Equations and parameters of the fit are indicated in Supplementary Information Table XI.

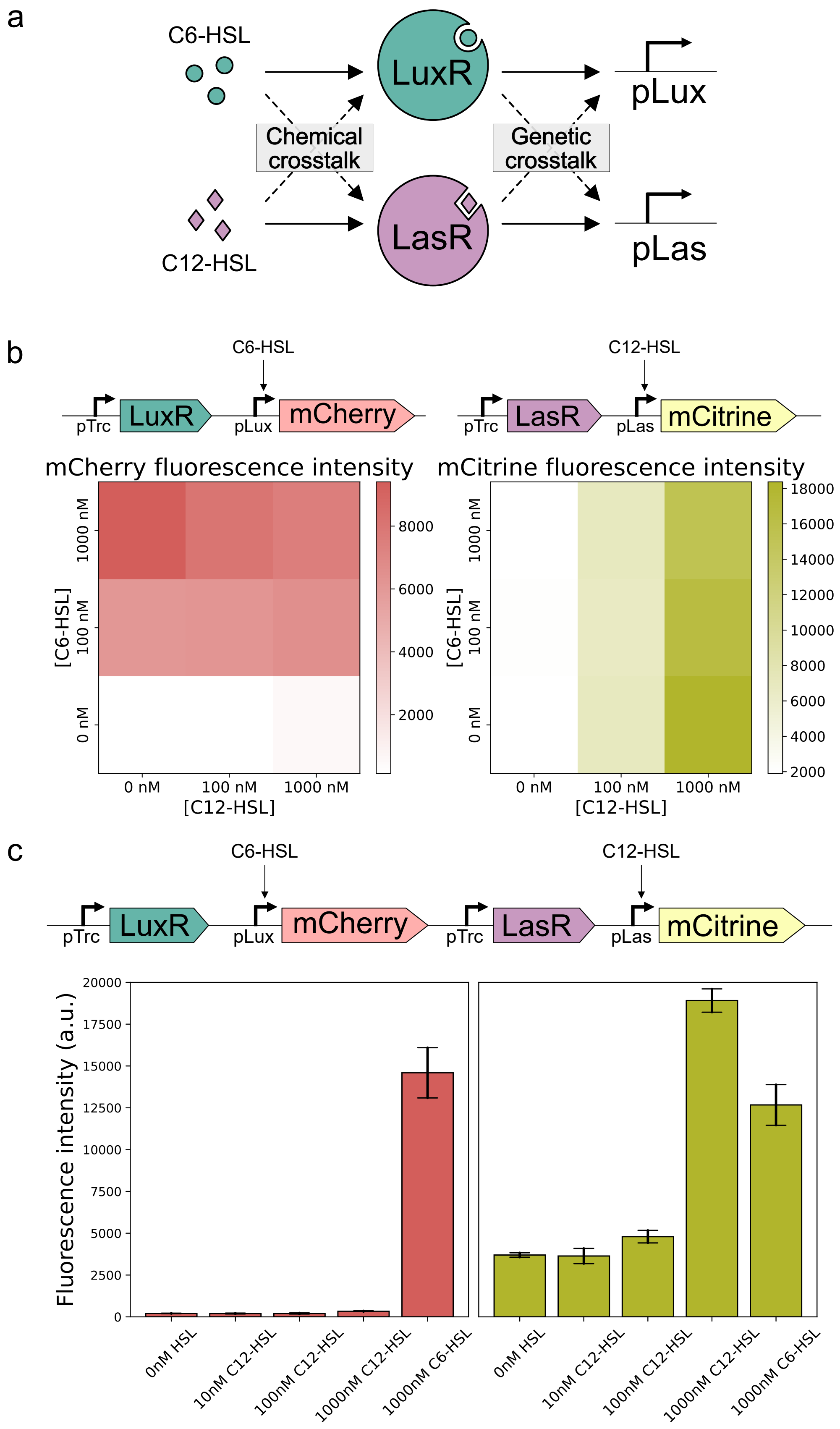
a
C6-HSL
LuxR
pLux
Chemical crosstalk
Genetic crosstalk
C12-HSL
LasR
pLas
b
C6-HSL
pTrc
LuxR
pLux
mCherry
C12-HSL
pTrc
LasR
pLas
mCitrine
mCherry fluorescence intensity
mCitrine fluorescence intensity
[C6-HSL]
1000 nM
100 nM
0 nM
[C12-HSL]
0 nM
100 nM
1000 nM
8000
6000
4000
2000
18000
16000
14000
12000
10000
8000
6000
4000
2000
c
C6-HSL
C12-HSL
pTrc
LuxR
pLux
mCherry
pTrc
LasR
pLas
mCitrine
Fluorescence intensity (a.u.)
20000
17500
15000
12500
10000
7500
5000
2500
0
0nM HSL
10nM C12-HSL
100nM C12-HSL
1000nM C12-HSL
1000nM C6-HSL
0nM HSL
10nM C12-HSL
100nM C12-HSL
1000nM C12-HSL
1000nM C6-HSL

Fig. S10. **Quantification of leakiness within, and cross talk between, the C6 and C12 devices.** a) Schematic representation of cross talk in the context of quorum sensing systems: chemical cross talk consists in a small diffusible molecule creating a functional complex with a non-cognate receptor; genetic crosstalk consists in a correct complex binding to, and activating, a non-cognate promoter. b) Fluorescence intensity of reporter cells carrying either the C6 (left side) or the C12 (right side) device (top panels, schematic). Cells were grown in a Biotek plate reader for 24 h in medium conditioned with different combinations of 3O-C6-HSL and 3O-C12-HSL, heatmaps represent the endpoint mCherry (red, left side) and mCitrine (yellow, right side) fluorescence intensity respectively. c) Fluorescence intensity of reporter cells carrying both the C6 and the C12 devices (top panel, schematic). Cells were grown in a Biotek plate reader for 24h in medium conditioned with different amounts of either 3O-C6-HSL or 3O-C12-HSL, bars represent the endpoint mCherry (red, left side) and mCitrine (yellow, right side) fluorescence intensity respectively (mean and standard error for 3 biological replicates). The C6 device showed minimal leakiness and cross talk, while the C12 device showed some degree of leakiness (around 20% of the maximal induction) and very strong cross talk (around 65% of the maximal induction).

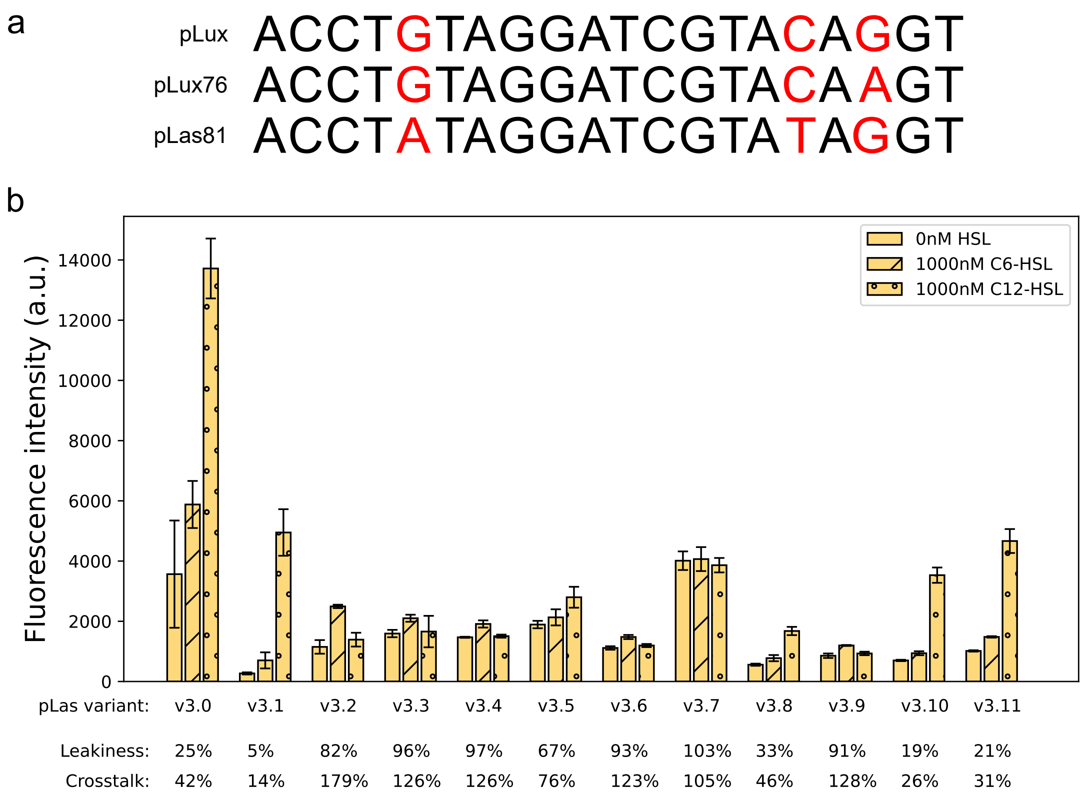


Fig. S11. **Screening of 12 pLas promoter variants.** a) The alignment of pLux, pLux76 and pLas81 promoter sequences showed a marked sequence identity. b) Quantification of leakiness and cross talk for each of the promoter variants listed in Supplementary Table III. Bars represent the mean and standard error (n = 2 biological replicates) of the endpoint mCitrine fluorescence intensity. For each variant, leakiness was calculated as the percentage of activity in absence of induction (empty bar) versus 3O-C12-HSL induction (dotted bar). Crosstalk was calculated as the percentage of activity with 3O-C6-HSL induction (striped bar) versus 3O-C12-HSL induction (dotted bar).

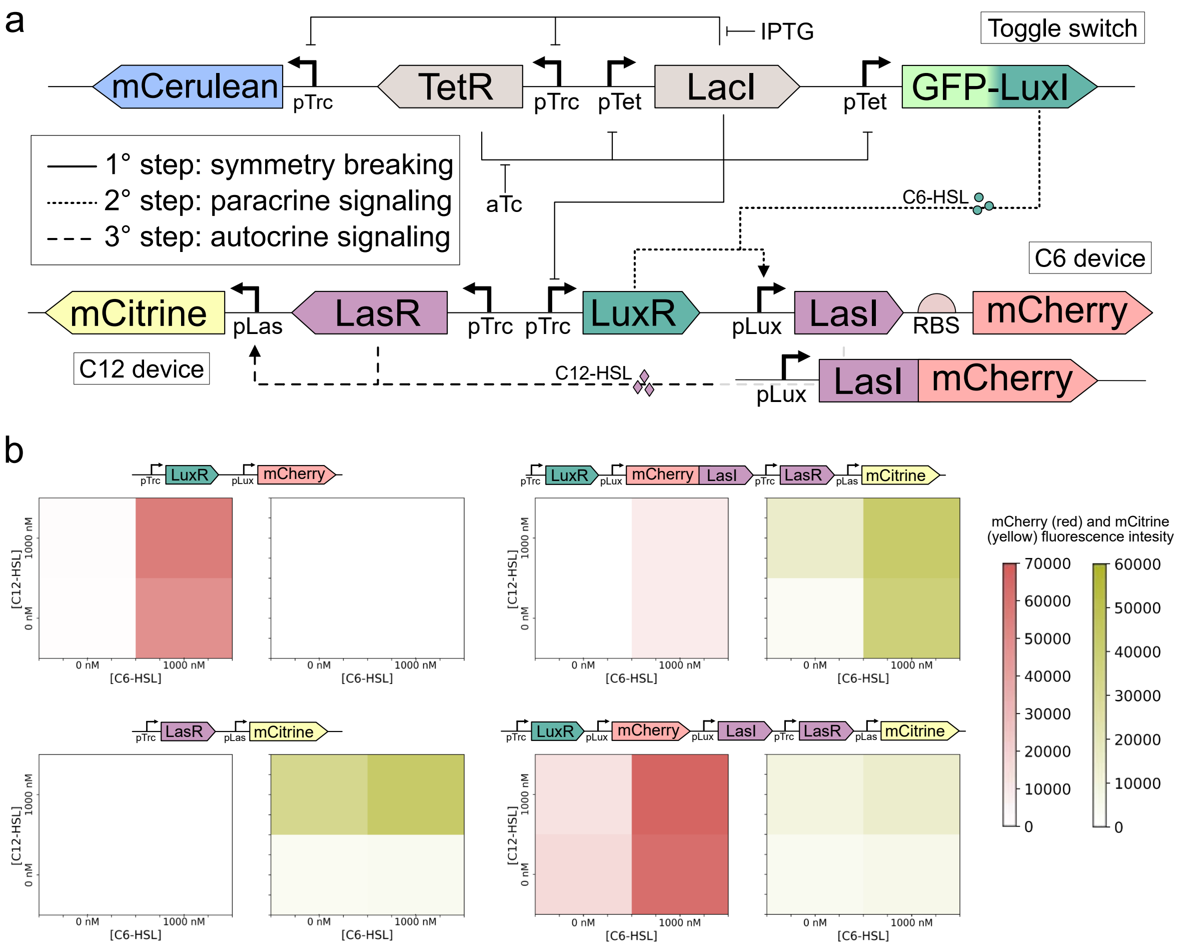


Fig. S12. **Attempts to engineer a 3-step differentiation system based on the quorum sensing systems LuxI-LuxR and LasI-LasR.** a) Genetic circuitry of the system, including interactions and chemical inducers. Coloured boxes: genes; L-shaped arrows: promoters; T-shaped lines: repressions; dashed and dashdot arrows: activations; green dots and purple diamonds: small diffusible molecules 3O-C6-HSL and 3O-C12-HSL, respectively. b) Fluorescence intensity of reporter cells carrying the C6 device (top left), the C12 device (bottom left), a fusion-protein version of the 3-step system (top right) and a bicistronic version of the 3-step system (bottom right). Top panels represent schematics of the circuits. Cells were grown in a Biotek plate reader for 24 h in medium conditioned with different combinations of 3O-C6-HSL and 3O-C12-HSL, heatmaps represent the endpoint mCherry (red, left side) and mCitrine (yellow, right side) fluorescence intensity respectively. Scale bars apply to all heatmaps.

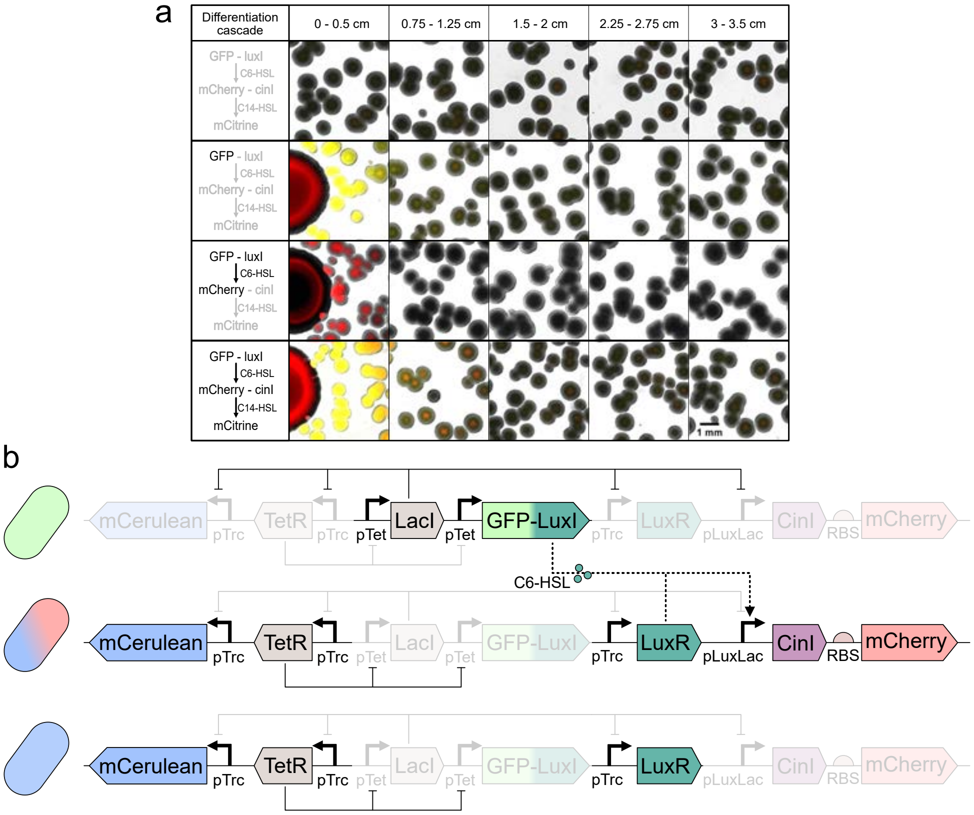


Fig. S13. **Optimization of the 3-step Lux-Cin system** a) Testing of the 3 sequential differentiation steps. The left panel shows the active (black) and inactive (gray) steps of the differentiation cascade. The right panel shows microscopy images of receiver colonies (small, on the right) grown in presence or absence of sender colonies (big, on the left), with complete or incomplete circuitry, in accordance with the left panel. I° row: receivers only. II° row: senders and receivers, the LuxI synthase is missing from the circuit. III° row: senders and receivers, the CinI synthase is missing from the circuit. IV° row: senders and receivers, complete circuity. Channels: brightfield, red, yellow. Leakiness from the pLux promoter led to mCherry expression and unwanted production of 3O-C14-HSL from green sender colonies, bypassing the $2^{nd}$ differentiation step. The sender colonies in this figure do not originate from one single cell, but from 1 $\mu$l of a culture of sender cells at OD=1, which explains their larger size (see Methods). b) Rationale for replacing pLux with pLuxLac. Schematic representation of the optimized 3-step circuit, with active (coloured) and inactive (transparent) parts depending on the cell identity (green, blue-red, blue, as marked by the stylized cell on the left). Coloured boxes: genes; L-shaped arrows: promoters; T-shaped lines: repressions; dashdot arrow: activation; green dots: small diffusible molecule 3O-C6-HSL. The pLuxLac promoter is tightly repressed by LacI in the green sender state, while it is available for the LuxR-C6 complex in the blue receiver state.

| Inducers in the plate | None | None | 1mM IPTG 1μM C6-HSL | None | None | 1mM IPTG 1μM C6-HSL |
|---|---|---|---|---|---|---|
| C6-HSL | − | 2 μl 0.5mM | − | − | − | − |
| C14-HSL | − | − | − | 2 μl 0.05mM | 2 μl 0.5mM | − |
| 3step SD system | − | − | − | − | − | + |

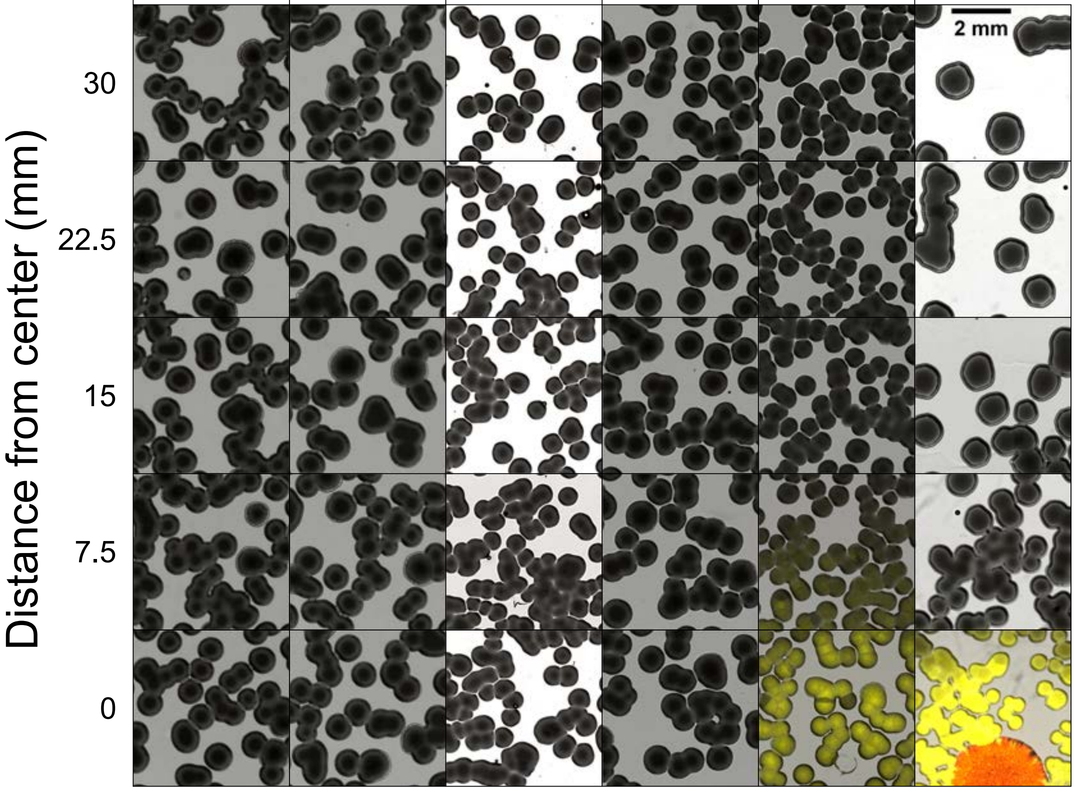


Fig. S14. **Characterization of the 3$^{rd}$ differentiation step in isolation.** Microscopy images of reporter colonies harbouring a 3O-C14-HSL responsive circuit (CinR_pCin-eYFP), grown in presence or absence of chemical inducers and the 3-step sequential differentiation system (combination marked in the top table). After 24 h incubation, 5 images were collected at increasing distances from the centre of the plate (distance marked on the left side of each row). A colony harbouring the 3-step sequential differentiation system and supplemented with 1 $\mu$M 3O-C6-HSL (6° column) resulted in 3O-C14-HSL production intermediate between 0.1 and 1 nano-moles of pure 3O-C14-HSL (4° and 5° column, respectively). The sender colony in the 6° column does not originate from one single cell, but from 1 $\mu$l of a culture of sender cells at OD=1, which explains its larger size (see Methods).

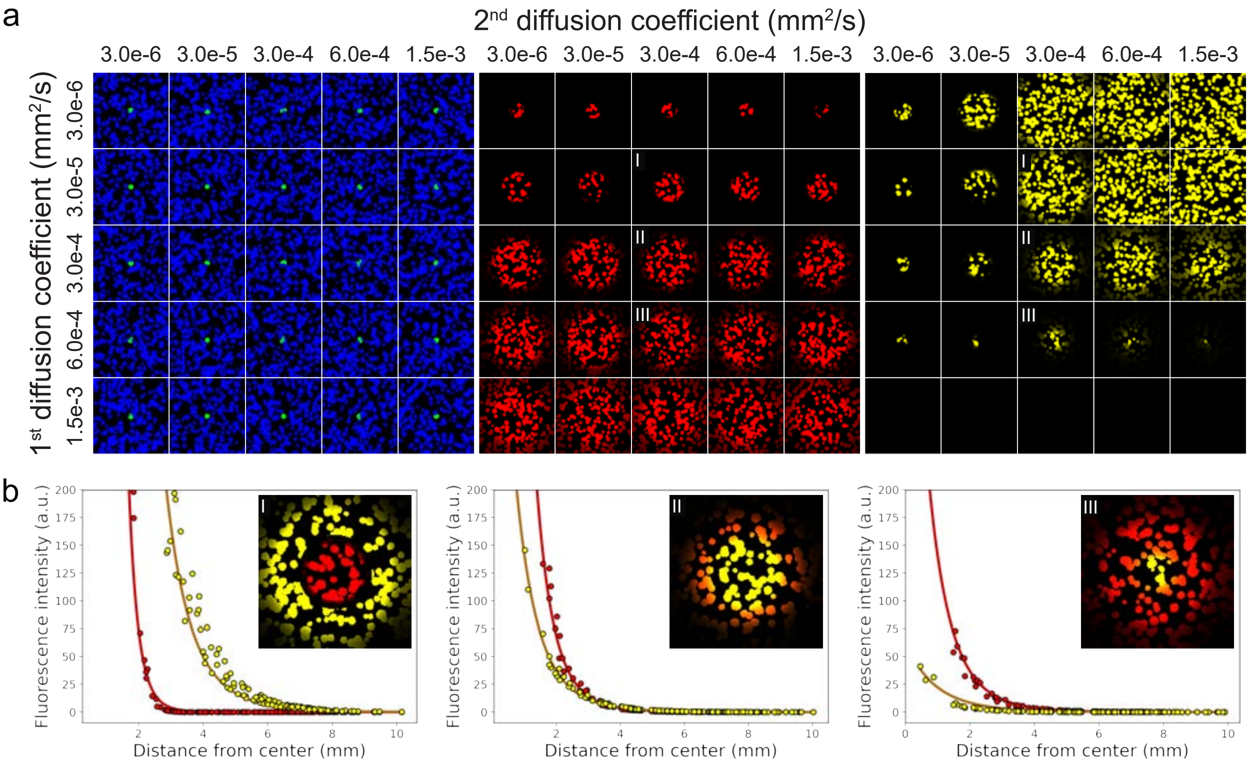


Fig. S15. **Effect of the diffusion coefficients on the spatial pattern generated from the 3-step differentiation system.** a) Mathematical simulations of the 3-step system were carried out varying diffusion coefficients for the two diffusible signals. Rows, from top to bottom: increasing diffusion coefficient for the $1^{st}$ signal. Columns, from left to right: increasing diffusion coefficient for the $2^{nd}$ signal. The default diffusion coefficient used for 3O-C6-HSL and 3O-C14-HSL is $3.0 \cdot 10^{-4}\ mm^2/s$ ($3^{rd}$ row, $3^{rd}$ column). Channels: left panel - green, blue; central panel - red; right panel - yellow. b) Quantification of the red and yellow fluorescence intensity of three simulations selected from a (marked with Roman numbers) show different profiles: I) a wide yellow ring surrounding a narrow red ring, II) roughly overlapping red and yellow rings, III) a wide red ring surrounding a narrow yellow ring. Points represent simulation points; black lines represent exponential curves fitted to simulation points. Equations and parameters of the fit are indicated in Supplementary Information Table XII.

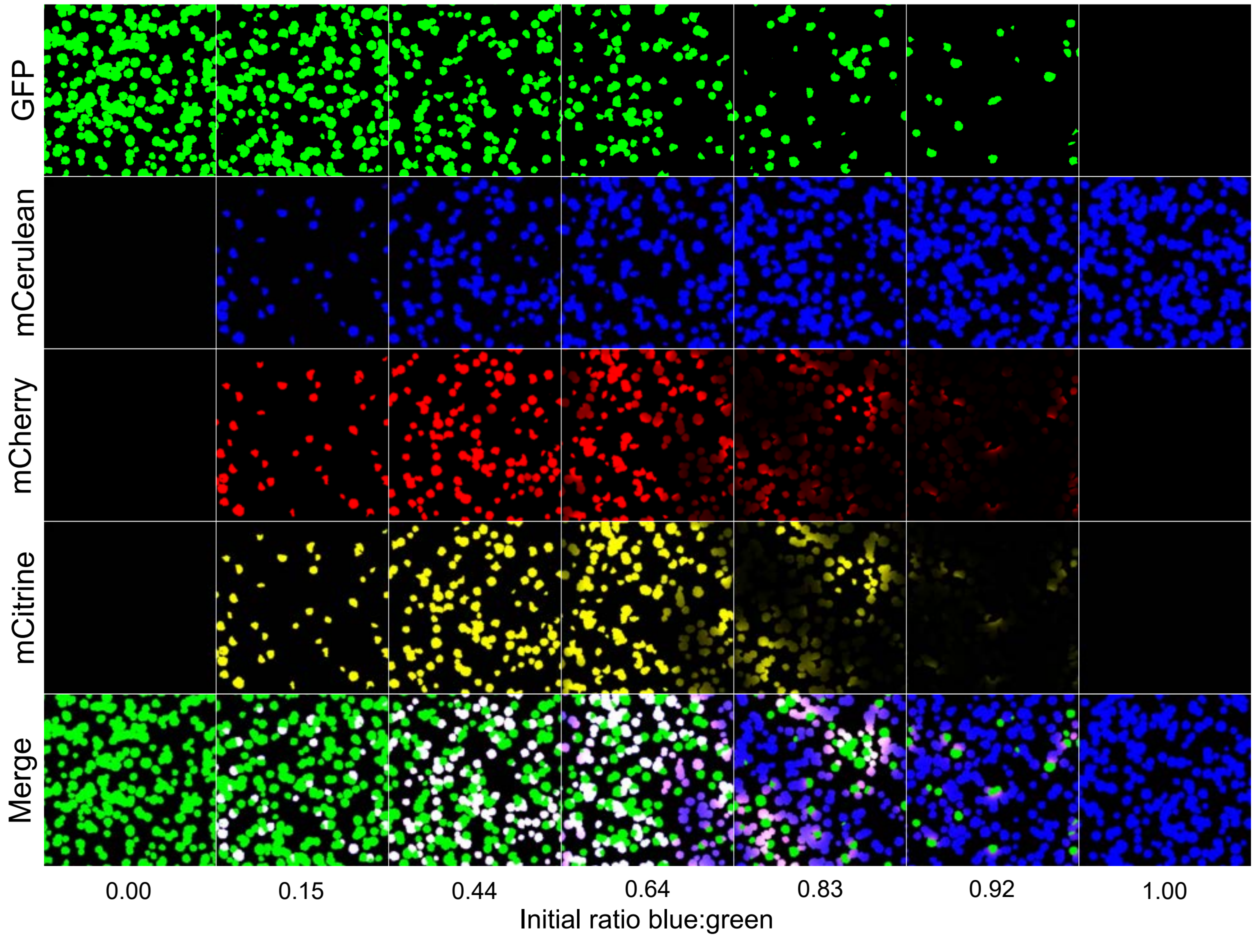


Fig. S16. **The 3-step sequential differentiation system can generate a variety of spatial patterns.** Mathematical simulation of spatial patterns at 24 h, starting from different blue:green ratios (indicated below each simulation). Colonies are colour-coded according to their simulated green (I° row), blue (II° row), red (III° row) and yellow (IV° row) fluorescence intensity. Initial blue:green ratios were chosen to match Figure 6d.

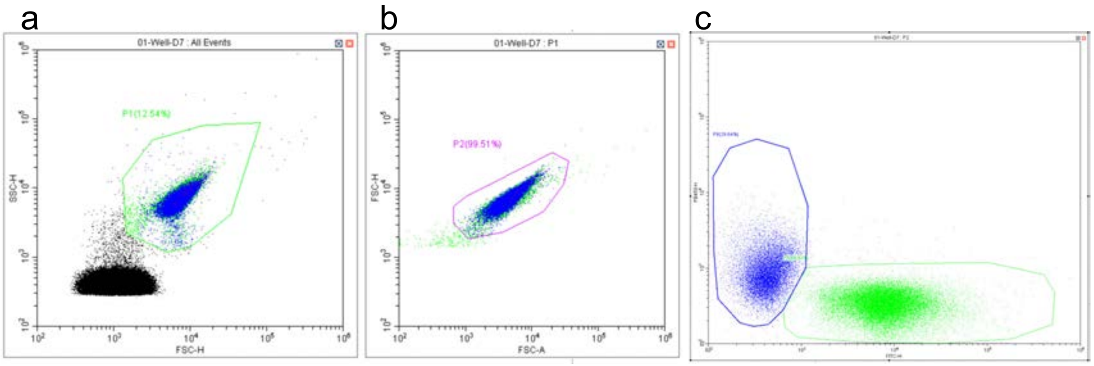


Fig. S17. **Gating procedure for Flow Cytometry data.** a) To discriminate between cells and other particles, all measured events were gated by forward scatter height (FSC.H) $> 10^3$ arbitrary unit (a.u.) and side scatter height (SSC.H) $> 10^3$ a.u, generating population P1 (green). b) We excluded doublets by plotting the FSC.H against the forward scatter area (FSC.A) and set a gate for the events with approximately 1:1 ratio of FSC.H to FSC.A, generating population P2 (purple). We recorded 50,000 events of singlet cells. c) To quantify the percentages of GFP-positive and mCerulean-positive cells we set gates below and above the y=x diagonal of the green (FITC-H) versus blue (PB450-H) graph, ensuring a signal of at least $10^2$ a.u. in the corresponding channel and at most $10^3$ a.u. in the opposite channel, generating populations P7 and P8 (green and blue, respectively).

### A. Mathematical modelling

#### 1. *Bacterial colonies growth*

The colony growth was described using a cellular automaton model [1]. In these models, the cell concentration ($c$) is normalized by the carrying capacity and binarized, such that the dimensionless concentration ($\hat{c}$) equals 1 when the surface is covered by cells, and 0 when it is cell-free. Thus, $\hat{c}$ represents the presence or absence of the bacterial colony. During the simulation, a free point of the computational grid (where $\hat{c} = 0$) becomes occupied ($\hat{c} = 1$) with a certain probability, capturing the expansion of the colony.

The probability $P(x, y)$ that an OFF grid point turns ON during a given time step is defined as

$$P(x, y) = p \left[ s(x, y) + \frac{m(x, y)}{\sqrt{2}} \right], \tag{1}$$

where $s(x, y)$ is the number of lateral and vertical neighbours in the $x$ and $y$ directions that are ON, and $m(x, y)$ is the number of diagonal (corner) neighbours that are ON. The parameter $p = 0.4$ was chosen to reproduce the experimentally observed colony shapes.

At the beginning of the simulation, which corresponds to 9h after colony seeding, a fixed number of points in the grid is turned ON. These points are distributed homogeneously on the grid, and are turned ON progressively during a fixed time interval, from 9h to 12h, until the desired colony density is matched, in order to recapitulate the experimentally observed variability in time of appearance and colony size. To match the colony growth dynamics observed in experiments, we performed time-lapse confocal microscopy measurements (Figure S19a,c). Using an intensity threshold below 600 (on a scale from 0 to 65535 where 0 and 65535 are the points with lowest and highest intensity, respectively), we quantified the fraction of the surface covered by the colonies. We then fitted the temporal growth curve with a double-sigmoid function, $\sigma_2(t)$, which captures growth driven by two distinct nutrient sources:

$$\sigma_2(t) = \frac{l_1}{1 + e^{-\lambda_1(t-t_1)}} + \frac{l_2}{1 + e^{-\lambda_2(t-t_2)}} \quad , \tag{2}$$

where $\lambda_1$ and $\lambda_2$ are the growth rate coefficients, $t_1$ and $t_2$ are the inflection points, and $l_1$ and $l_2$ represent the maximum contributions associated with the two nutrient regimes. An

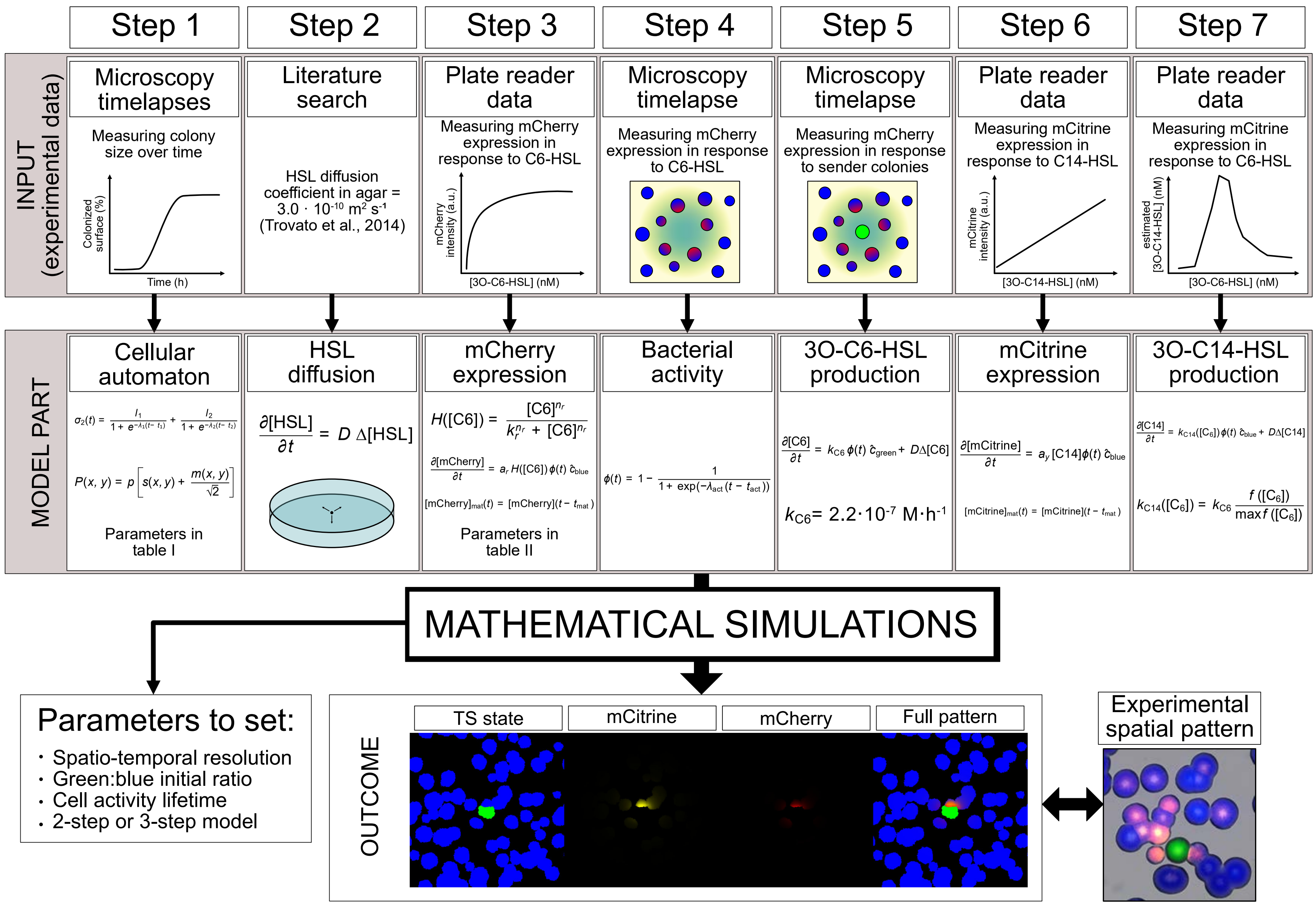

Step 1
Step 2
Step 3
Step 4
Step 5
Step 6
Step 7
INPUT (experimental data)
Microscopy timelapses
Measuring colony size over time
Colonized surface (%)
Time (h)
Literature search
HSL diffusion coefficient in agar = $3.0 \cdot 10^{-10}$ $m^2$ $s^{-1}$ (Trovato et al., 2014)
Plate reader data
Measuring mCherry expression in response to C6-HSL
mCherry intensity (a.u.)
[3O-C6-HSL] (nM)
Microscopy timelapse
Measuring mCherry expression in response to C6-HSL
Microscopy timelapse
Measuring mCherry expression in response to sender colonies
Plate reader data
Measuring mCitrine expression in response to C14-HSL
mCitrine intensity (a.u.)
[3O-C14-HSL] (nM)
Plate reader data
Measuring mCitrine expression in response to C6-HSL
estimated [3O-C14-HSL] (nM)
[3O-C6-HSL] (nM)
MODEL PART
Cellular automaton
$\sigma_2(t) = \frac{l_1}{1+e^{-\lambda_1(t-t_1)}} + \frac{l_2}{1+e^{-\lambda_2(t-t_2)}}$
$P(x,y) = p\left[s(x,y) + \frac{m(x,y)}{\sqrt{2}}\right]$
Parameters in table I
HSL diffusion
$\frac{\partial[HSL]}{\partial t} = D\,\Delta[HSL]$
mCherry expression
$H([C6]) = \frac{[C6]^{n_r}}{k_r^{n_r} + [C6]^{n_r}}$
$\frac{\partial[mCherry]}{\partial t} = a_r H([C6])\,\phi(t)\,\hat{c}_{blue}$
$[mCherry]_{mat}(t) = [mCherry](t - t_{mat})$
Parameters in table II
Bacterial activity
$\phi(t) = 1 - \frac{1}{1+\exp(-\lambda_{act}(t - t_{act}))}$
3O-C6-HSL production
$\frac{\partial[C6]}{\partial t} = k_{C6}\,\phi(t)\,\hat{c}_{green} + D\Delta[C6]$
$k_{C6} = 2.2 \cdot 10^{-7}$ $M \cdot h^{-1}$
mCitrine expression
$\frac{\partial[mCitrine]}{\partial t} = a_y [C14]\,\phi(t)\,\hat{c}_{blue}$
$[mCitrine]_{mat}(t) = [mCitrine](t - t_{mat})$
3O-C14-HSL production
$\frac{\partial[C14]}{\partial t} = k_{C14}([C_6])\,\phi(t)\,\hat{c}_{blue} + D\Delta[C14]$
$k_{C14}([C_6]) = k_{C6}\,\frac{f([C_6])}{\max f([C_6])}$
MATHEMATICAL SIMULATIONS
Parameters to set:
· Spatio-temporal resolution
· Green:blue initial ratio
· Cell activity lifetime
· 2-step or 3-step model
OUTCOME
TS state
mCitrine
mCherry
Full pattern
Experimental spatial pattern

Fig. S18. **Overview of the simulation workflow. Step 1)** Colony expansion was modelled using cellular automaton simulations to match experimentally observed growth dynamics. **Step 2)** Diffusion of signaling molecules in the agar was computed in three dimensions using literature-derived diffusion coefficients. **Step 3)** mCherry production rate was quantified from plate-reader measurements where receiver cells were grown in presence of 3O-C6-HSL. **Step 4)** Bacterial activity was quantified from microscopy time-lapses where receiver colonies were grown in presence of 3O-C6-HSL. **Step 5)** 3O-C6-HSL production rate was quantified from microscopy time-lapses where receiver colonies were grown in presence of sender cells. **Step 6)** mCitrine production rate was quantified from plate-reader measurements where receiver cells were grown in presence of 3O-C14-HSL. **Step 7)** By relating mCitrine levels to 3O-C6-HSL concentration, we estimated 3O-C14-HSL production. These experimentally constrained parameters together informed the spatial simulations, which were then compared to experimental observations. Details of each parametrization step are provided in the SI Sections A1-7 (section numbers corresponding to step numbers in the figure).

alternative explanation for the observed change in growth rate is a metabolic switch, whereby cells transition between different metabolic pathways as the initially available substrate becomes depleted [2]. The fitted parameters are summarized in Table I. At each time step, new cells were added to the colony in the cellular automaton model until the simulated growth matched the experimentally determined curve (Figure S19b,d). For simplicity, the same growth strategy was applied to all cell states (green, blue . . . ), assuming that the growth rate is independent of the cellular state.

TABLE I. Fitted parameters of the double-sigmoid function (Eq. 1) used to describe the experimentally observed growth rates in Figure S19a.

| $l_1$ | $l_2$ | $\lambda_1$ | $\lambda_2$ | $t_1$ | $t_2$ |
|---|---|---|---|---|---|
| 0.1358 | 0.2583 | 1.9173 $h^{-1}$ | 1.1635 $h^{-1}$ | 12.21 h | 14.81 h |

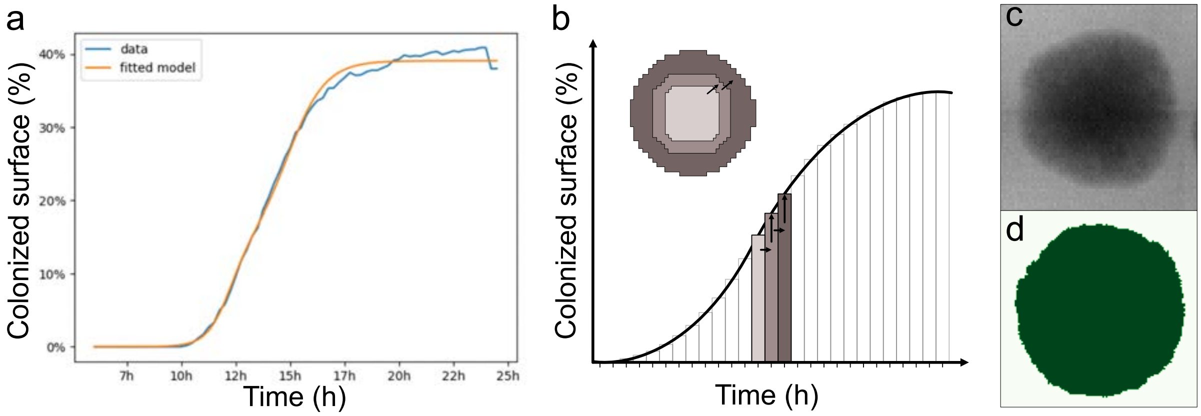


Fig. S19. **Parametrization and functioning of the cellular automaton framework.** a) Experimental colony expansion quantified as the percentage of surface area covered over time, together with the fitted double-sigmoid growth curve (Eq. 1). b) Schematic representation of the cellular automaton functioning: for each time-step (right-facing arrows) the cellular automaton performs as many expansion steps (up-facing arrows) as needed for the simulated colony to match the experimentally observed colony expansion. The insert shows a visualization of the colony expansion over two time-steps. c) Brightfield image of an experimentally grown colony after 24 h incubation. d) Colony generated using the cellular automaton model for a time period of 24 h.

#### *2. Diffusion of HSLs*

The HSLs (3O-C6-HSL and 3O-C14-HSL) produced by sender cells diffuse through the agar according to the diffusion equation:

$$\frac{\partial[\mathrm{HSL}]}{\partial t} = D\,\Delta[\mathrm{HSL}] \quad , \tag{3}$$

where [HSL] is the concentration of HSL, $D$ is the diffusion coefficient in agar, set to $3 \cdot 10^{-10}$ m$^2$/s for both C6-HSL and C14-HSL [3], and $\Delta$ is the Laplacian operator. This equation was solved on a three-dimensional domain, as the depth of the agar is non-negligible relative to the diffusion timescale of HSLs during the biochemical processes.

We solved the diffusion equation on a uniform grid using a finite-difference scheme with explicit Euler integration [4], employing a spatial step of 0.1 mm and a time step of 2.4 s. Periodic boundary conditions were applied in the $x$ and $y$ directions, and no-flux boundary conditions in the $z$ direction. Initially, the domain contained no HSLs; they were produced at the surface by the cells as described in SI Sections A 5 and A 7.

#### 3. *mCherry production and 3O-C6-HSL sensing*

mCherry is produced by cells in the blue receiver state when they sense 3O-C6-HSL. However, cellular activity declines over time, which we accounted for using an activity function $\phi(t)$ (see next section). We also assumed that the 3O-C6-HSL response is non-linear and follows a Hill function $\mathcal{H}$. Under these assumptions, the mCherry concentration ([mCherry]) evolves according to

$$\frac{\partial[\mathrm{mCherry}]}{\partial t} = a_r\,\mathcal{H}([\mathrm{C6}])\,\phi(t)\,\hat{c}_{\mathrm{blue}} \quad , \tag{4}$$

where $a_r$ is the mCherry production rate coefficient and $\hat{c}_{\mathrm{blue}}$ is the concentration of cells in the blue state in the cellular automaton model. The Hill response to 3O-C6-HSL is given by

$$\mathcal{H}([\mathrm{C6}]) = \frac{[\mathrm{C6}]^{n_r}}{k_r^{n_r} + [\mathrm{C6}]^{n_r}} \quad , \tag{5}$$

where $k_r$ and $n_r$ are, respectively, the half-saturation constant and Hill coefficient for mCherry induction by 3O-C6-HSL. Furthermore, we assumed that the proteins undergo a maturation phase, and become detectable only after a maturation time $t_{\mathrm{mat}}$:

$$[\mathrm{mCherry}]_{\mathrm{mat}}(t) = [\mathrm{mCherry}](t - t_{\mathrm{mat}}) \quad . \tag{6}$$

To determine the parameters $k_r$ and $n_r$ of the mCherry production model, we first used plate reader measurements. In liquid media, we measured red fluorescence at various 3O-C6-HSL concentrations after 48 hours (Figure S3 for the 2-step system and Figure S20 for the 3-step system) and fitted a Hill function (Eq. 5) to the data. The parameters fitted for mCherry production in the 2-step and 3-step systems are quite different. We hypothesize that the following elements contribute to the difference: a) the replacement of pLux76 with pLuxLac, which leads to tighter regulation and overall reduced expression of the downstream genes; b) the addition of the cinI synthase and a RBS in a monocistronic unit upstream of mCherry, which might justify reduced expression of mCherry; c) circuit-induced cell burden and reduced growth rate for the 3-step system, which leads to a slower response and lower reporter expression.

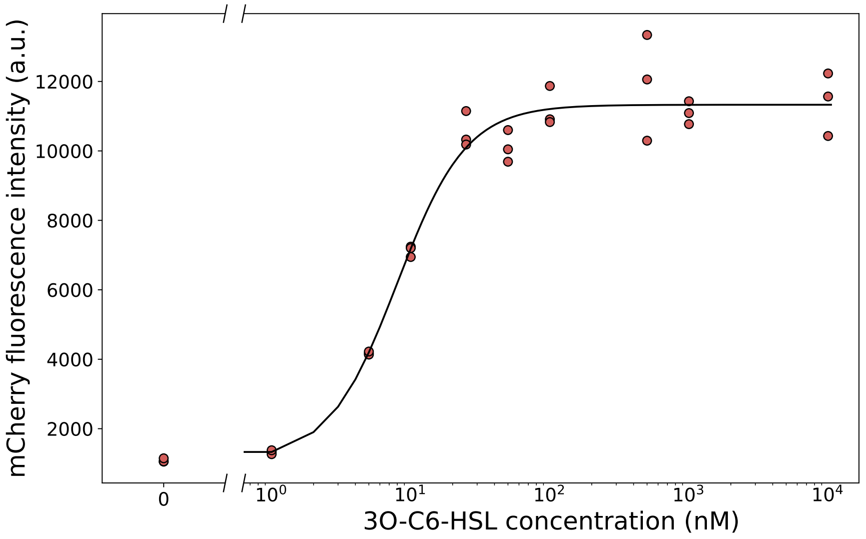


Fig. S20. **Characterization of mCherry fluorescence intensity as a function of 3O-C6-HSL concentration in the 3-step system.** Cells containing the 3-step system (toggle switch, C6 device and C14 device) were grown in a Biotek plate reader for 24 h in medium conditioned with different concentrations of 3O-C6-HSL. Points represent the end-point red fluorescence intensity of 3 biological replicates per condition; black line represents a Hill function (Eq. 5) fitted to experimental points (Equation: $y = \frac{a}{b+x^c} + d$. Parameters: $a$ = 270.94, $b$ = 0.264, $c$ = -1.747, $d$ = 1065.885. Method: Levenberg-Marquardt algorithm).

#### *4. Bacterial activity*

In our system, production of diffusible molecules and fluorescent reporters happens on a timescale of several hours, therefore we decided to take into account the entry of cells in stationary phase. Traditionally, bacterial activity has been modelled as a step-wise inactivation function [5]. However, experimental evidence has shown that bacteria support a low constant rate of protein expression even while growth-arrested, suggesting a low decay in bacterial activity [6]. Even when considering cell death upon heat inactivation, a Weibull frequency distribution model is preferred over a step-wise viability function [7, 8]. While we could not find mathematical descriptions specifically for entry in stationary phase, several single-cell metabolism studies support gradual, asynchronous state transitions [9–11]. We therefore decided to describe the loss of activity function in our system via a monotonically decreasing sigmoid function, that captured well our experimental data.

The bacterial activity function is given by the sigmoid function

$$\phi(t) = 1 - \frac{1}{1 + \exp(-\lambda_{\mathrm{act}}(t - t_{\mathrm{act}}))} \quad , \tag{7}$$

where $\lambda_{\mathrm{act}}$ and $t_{\mathrm{act}}$ describe the onset and steepness of the activity decline, respectively.

To complete the fitting of all parameters required for the mCherry production model, we performed time-lapse experiments on solid surface. Receiver colonies were grown in presence of a droplet of 2 $\mu$l of 3O-C6-HSL 50 $\mu$M which could diffuse in the medium (Supplementary Movie 1). Building upon a simplified version of the mCherry production model, we fine-tuned the two parameters for the activity function, $\lambda_{act}$ and $t_{act}$, and the mCherry maturation time $t_{\mathrm{mat}}$. Our manually fitted mCherry maturation time was consistent with values previously reported in the literature [12]. All fitted parameters for the mCherry production model are summarized in Table II. For the implementation of our mathematical model, we were not interested in analysing individual parameters values, but rather in the resulting response function to varying 3O-C6-HSL concentrations. Further analysis would be needed to determine whether these parameters are statistically significant, but this is beyond the scope of this present manuscript.

TABLE II. Parameters used for mCherry production. The parameters $a_r$, $n_r$, and $k_r$ were obtained by fitting the data shown in Figure S3, $\lambda_{act}$, $t_{act}$ and $t_{mat}$ were manually fine-tuned. The first and second rows correspond to parameter sets without and with 30-C14-HSL and mCitrine production, respectively.

| $a_r$ (a.u./h) | $n_r$ | $k_r$ (nM) | $\lambda_{act}$ | $t_{act}$ (h) | $t_{mat}$ (h) |
|---|---|---|---|---|---|
| $1.489 \cdot 10^4$ | 0.8397 | 242.4 | 2 | 18.5 | 1 |
| $1.026 \cdot 10^4$ | 1.747 | 37.88 | 2 | 18.5 | 1 |

#### *5. 3O-C6-HSL production*

To determine the 3O-C6-HSL production rate of green sender cells, we performed a time-lapse experiment similar to that described in the previous section, where the 3O-C6-

HSL droplet was replaced by sender cells (Supplementary Movie 3). Receiver cells with previously characterized parameters acted as biosensors, enabling us to calculate the 3O-C6-HSL production rate from green colonies. We assumed that the 3O-C6-HSL production is proportional to the surface coverage of sender cells, $\hat{c}_{\mathrm{green}}$, and their biological activity, $\phi(t)$:

$$\frac{\partial[\mathrm{C6}]}{\partial t} = k_{\mathrm{C6}}\,\phi(t)\,\hat{c}_{\mathrm{green}} + D\Delta[\mathrm{C6}] \quad , \tag{8}$$

where $D$ is the diffusion coefficient and $k_{\mathrm{C6}}$ is the 3O-C6-HSL production rate coefficient. Based on our experiments, the fitted value is $k_{\mathrm{C6}} = 2.2 \cdot 10^{-7}\ \mathrm{M} \cdot \mathrm{h}^{-1}$.

#### 6. *mCitrine production and 3O-C14-HSL sensing*

When extending the experimental system with mCitrine and 3O-C14-HSL production, we found that the mCherry production parameters also changed. Thus, for the full system including mCitrine, we used the parameter set from the second row of Table II to describe mCherry production. To determine the mCitrine sensing capability, we applied the same strategy used for mCherry: plate reader measurements were performed across a range of 3O-C14-HSL concentrations. Although one would expect the yellow fluorescence to saturate at high 3O-C14-HSL levels —yielding a Hill-type response analogous to mCherry— we observed a linear response over the biologically relevant concentration range (Figure S21a). Motivated by this, we assumed that mCitrine production is linearly proportional to the 3O-C14-HSL concentration [C14]:

$$\frac{\partial[\mathrm{mCitrine}]}{\partial t} = a_y\,[\mathrm{C14}]\,\phi(t)\,\hat{c}_{\mathrm{blue}} \quad , \tag{9}$$

where $\phi(t)$ and $\hat{c}_{\mathrm{blue}}$ represent the biological activity (from section A 4) and the concentration of cells in the blue state in the cellular automaton model, and $a_y = 1$ a.u./(h $\cdot$ nM) is the mCitrine production rate coefficient. When plotting the yellow fluorescence intensity, we incorporated a maturation delay for mCitrine, using the same formulation applied to mCherry in Eq. 6, and $t_{\mathrm{mat}} = 15\ min$:

$$[\mathrm{mCitrine}]_{\mathrm{mat}}(t) = [\mathrm{mCitrine}](t - t_{\mathrm{mat}}) \quad . \tag{10}$$

Our manually fitted maturation time for mCitrine was similar to the experimentally measured maturation time of the closest fluorophore relative, EYFP [13].

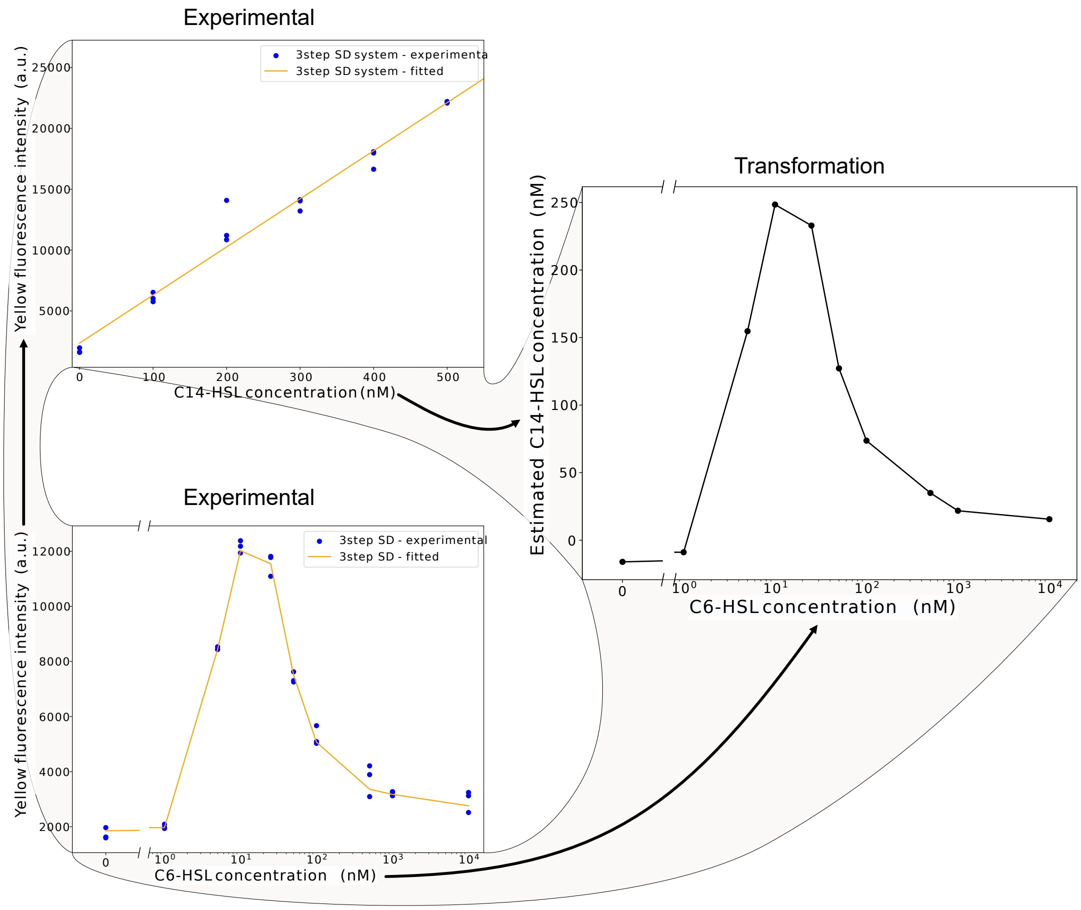


Fig. S21. **Estimating the 3O-C14-HSL production rate from plate reader measurements.** a) Yellow fluorescence intensity as a function of externally supplied 3O-C14-HSL in receiver cells. Blue dots represent experimental points (3 biological replicates per condition), yellow line represents a linear fit (slope = 39.51, intercept = 2357.47). b) Yellow fluorescence intensity as a function of 3O-C6-HSL, where 3O-C6-HSL induces 3O-C14-HSL production in receiver cells, which in turn activates mCitrine expression. Blue dots represent experimental points (3 biological replicates per condition), yellow line represents the response (a broken line connecting the means of the experimental points). c) Estimated 3O-C14-HSL concentration as a function of 3O-C6-HSL, obtained by combining the calibration in panel a with the C6–fluorescence relationship in panel b. mCitrine fluorescence intensity was quantified at 24 hours.

*7. 3O-C14-HSL production*

Finally, we estimated the 3O-C14-HSL production rate of receiver cells in the red state. To this end, we performed plate-reader experiments in which we measured the yellow fluorescence intensity as a function of 3O-C6-HSL concentration. In the presence of 3O-C6-HSL, receiver cells switch to the red state and begin producing 3O-C14-HSL; this 3O-C14-HSL, in turn, induces mCitrine production. We quantified the resulting mCitrine fluorescence after 24 hours (Figure S21b). Using the previously measured C14–yellow intensity relationship, we converted this curve into a C14–C6 response function (Figure S21c), which describes how much 3O-C14-HSL is produced at a given 3O-C6-HSL level. We assumed that the ratio of production rates, $k_{\mathrm{C14}}/k_{\mathrm{C6}}$, is proportional to this response function and normalized such that the proportionality is equal to one at its maximum. The 3O-C14-HSL production rate at a given 3O-C6-HSL concentration was therefore estimated by interpolation:

$$k_{\mathrm{C14}}([\mathrm{C}_6]) = k_{\mathrm{C6}} \frac{f([\mathrm{C}_6])}{\max f([\mathrm{C}_6])} \quad , \tag{11}$$

where $k_{\mathrm{C14}}$ and $k_{\mathrm{C6}}$ are the 3O-C14-HSL and 3O-C6-HSL production rates, respectively, and $f$ denotes the experimentally determined C14–C6 relationship. The 3O-C14-HSL concentration dynamics were then computed analogously to 3O-C6-HSL (Eq. 8):

$$\frac{\partial[\mathrm{C14}]}{\partial t} = k_{\mathrm{C14}}([\mathrm{C}_6])\, \phi(t)\, \hat{c}_{\mathrm{blue}} + D\Delta[\mathrm{C14}] \quad , \tag{12}$$

where $\hat{c}_{\mathrm{blue}}$ is the concentration of cells in the blue state in the cellular automaton model, and the remaining parameters are identical to those in Eq. 8.

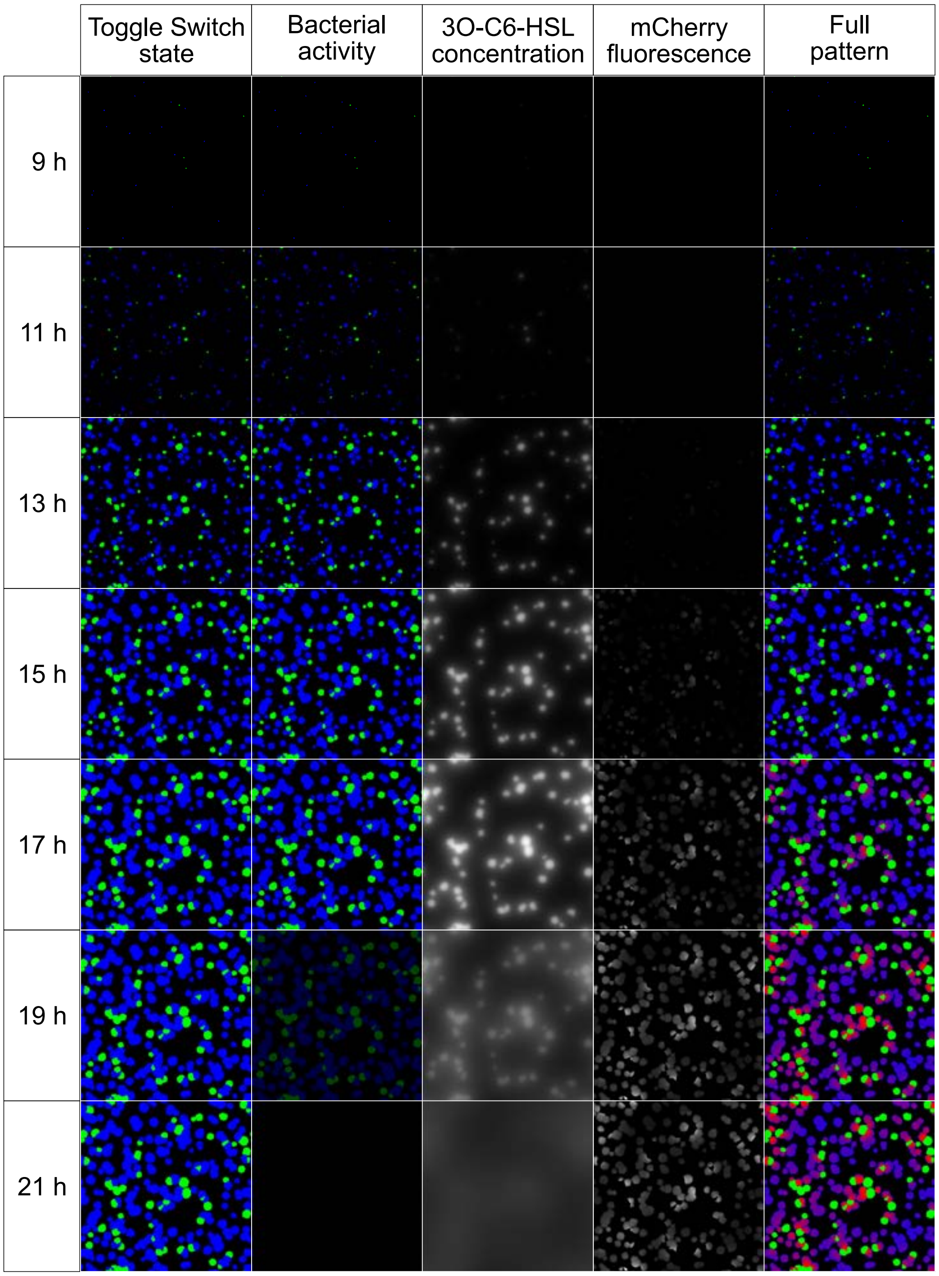


Fig. S22. **A simulation of the 2-step differentiation system**. The element visualized in each panel is marked on top of the corresponding column. Simulated time from the beginning of the experiment is marked on the left of the corresponding row.

TABLE III. **List of pLas promoter variants.**

| pLas variant | Name | Reference |
|---|---|---|
| v3.0 | pLas81 | Grant et al. [14] |
| v3.1 | pLas_iGem_short | iGem distribution kit 2024 |
| v3.2 | pLas82 | Grant et al. [14] |
| v3.3 | plasO-J23106 | Tica et al. [1] |
| v3.4 | NonCooperativeConsensus_short | Gilbert et al. [15] |
| v3.5 | NonCooperativeConsensus_long | Gilbert et al. [15] |
| v3.6 | CooperativeConsensus_short | Gilbert et al. [15] |
| v3.7 | pJ23106-lasO | Tica et al. [1] |
| v3.8 | Native_pLas_short | iGem distribution kit 2024 |
| v3.9 | CooperativeConsensus_long | Gilbert et al. [15] |
| v3.10 | pLas81_short | Curatolo et al. [16] |
| v3.11 | pLas81_long | Curatolo et al. [16] |

TABLE IV. **Plasmids used in this work.**

| **Plasmid** | **Features** | **Resistance** | **Origin** | **Addgene ID** |
|---|---|---|---|---|
| TS | Toggle switch (TetR - mCerulean / LacI - GFP) | Kam | ColE1 | 251146 |
| TS-LuxI | Toggle switch (TetR - mCerulean / LacI - GFP) with GFP-LuxI fusion protein | Kam | ColE1 | 251147 |
| C6_device v0 | LuxR receptor + pLux-mCherry | Spec | CDF | |
| C6_device v1 | LuxR receptor + pLux54%-mCherry | Spec | CDF | |
| C6_device v2 | LuxR receptor + pLux54%-G1-mCherry | Spec | CDF | |
| C6_device v3 | LuxR receptor + pLuxG1-mCherry | Spec | CDF | |
| C6_device v4 | LuxR receptor + pLux76-mCherry | Spec | CDF | 251148 |
| C6_device v5 | LuxR receptor + pLux54%-76-mCherry | Spec | CDF | |
| C12_device v3.0 | LasR receptor + pLas v3.0-mCitrine | Spec | CDF | |
| C12_device v3.1 | LasR receptor + pLas v3.1-mCitrine | Spec | CDF | |
| C12_device v3.2 | LasR receptor + pLas v3.2-mCitrine | Spec | CDF | |
| C12_device v3.3 | LasR receptor + pLas v3.3-mCitrine | Spec | CDF | |
| C12_device v3.4 | LasR receptor + pLas v3.4-mCitrine | Spec | CDF | |
| C12_device v3.5 | LasR receptor + pLas v3.5-mCitrine | Spec | CDF | |
| C12_device v3.6 | LasR receptor + pLas v3.6-mCitrine | Spec | CDF | |
| C12_device v3.7 | LasR receptor + pLas v3.7-mCitrine | Spec | CDF | |
| C12_device v3.8 | LasR receptor + pLas v3.8-mCitrine | Spec | CDF | |
| C12_device v3.9 | LasR receptor + pLas v3.9-mCitrine | Spec | CDF | |
| C12_device v3.10 | LasR receptor + pLas v3.10-mCitrine | Spec | CDF | |
| C12_device v3.11 | LasR receptor + pLas v3.11-mCitrine | Spec | CDF | |
| C6_C12_device v0 | LuxR receptor + pLux-mCherry, LasR receptor + pLas-mCitrine | Spec | CDF | |
| C6_C12_device v1 | LuxR receptor + pLux-mCherry-LasI, LasR receptor + pLas-mCitrine | Spec | CDF | |
| C6_C12_device v2 | LuxR receptor + pLux-mCherry + pLux-LasI, LasR receptor + pLas-mCitrine | Spec | CDF | |
| C14_device | CinR receptor + pCin-eYFP | Kam | ColA | 251149 |
| C6_C14_device v0 | LuxR receptor + pLux-CinI-mCherry, CinR receptor + pCin-mCitrine | Spec | CDF | |
| C6_C14_device v1 | LuxR receptor + pLuxLac-CinI-mCherry, CinR receptor + pCin-mCitrine | Spec | CDF | 251150 |

TABLE V. **Transformed strains used in this work.** The host strain is always *E. coli* MG1655

| **Strain name** | **Plasmid 1** | **Plasmid 2** | **Figure(s)** |
|---|---|---|---|
| 1-step system | TS-LuxI | | 2b-e, 4, S1, S2 |
| C6 reporter v0 | C6_device v0 | | S4 |
| C6 reporter v1 | C6_device v1 | | S4 |
| C6 reporter v2 | C6_device v2 | | S4 |
| C6 reporter v3 | C6_device v3 | | S4 |
| C6 reporter v4 | C6_device v4 | | 4, S3, S4, S8, S10b, S12b |
| C6 reporter v5 | C6_device v5 | | S4 |
| 2-step system | TS-LuxI | C6_device v4 | 5a-b, 6b-c, 7c, S5, S6, S7, S8, S13a, S17, S18, S19 |
| C12 reporter v0 | C12_device v3.0 | | S10b, S11, S12b |
| C12 reporter v1 | C12_device v3.1 | | S11 |
| C12 reporter v2 | C12_device v3.2 | | S11 |
| C12 reporter v3 | C12_device v3.3 | | S11 |
| C12 reporter v4 | C12_device v3.4 | | S11 |
| C12 reporter v5 | C12_device v3.5 | | S11 |
| C12 reporter v6 | C12_device v3.6 | | S11 |
| C12 reporter v7 | C12_device v3.7 | | S11 |
| C12 reporter v8 | C12_device v3.8 | | S11 |
| C12 reporter v9 | C12_device v3.9 | | S11 |
| C12 reporter v10 | C12_device v3.10 | | S11 |
| C12 reporter v11 | C12_device v3.11 | | S11 |
| C6-C12 reporter | C6_C12_device v0 | | S10c |
| 3-step Lux-Las system v1 | TS-LuxI | C6_C12_device v1 | S12b |
| 3-step Lux-Las system v2 | TS-LuxI | C6_C12_device v2 | S12b |
| C14 reporter | C14_device | | S14 |
| 3-step Lux-Cin system v0 | TS-LuxI | C6_C14_device v0 | S13a |
| 3-step Lux-Cin system v1 | TS-LuxI | C6_C14_device v1 | 1, 7c-e, S14, S20, S21 |
| Incomplete cascade v0 | TS | C6_C14_device v0 | S13a |
| Incomplete cascade v1 | TS | C6_C14_device v1 | 7c |

### TABLE VI. **List of primers used in this work.**

| Name | Sequence 5' - 3' | Sense | Description |
|---|---|---|---|
| PR_EB_161 | tcgctgggacgcccgagtattaactatcgttcaactg | F | Replace pLux promoter - binding site: suffix + RBS KDL027 |
| PR 419 | cagcctgcggtccggacctgtaggatcgtacatgctgacattgtgagcggataacaatactgtttcttcaatcgctgggacgcccg | F | Replace pLux promoter - Plux54%: promoter with 54% strength compared to the original pLux |
| HK524 | cagcctgcggtccggaccagtaggatcgtagaggtctgacattgtgagcggataacaatactgtttcttcaatcgctgggacgcccg | F | Replace pLux promoter - Plux54%-G1: lux box with lower affinity for LuxR, promoter with 54% strength compared to the original pLux |
| PR_EB_165 | ccggaccgcaggctgtccaagcgattgctgaggtc | R | Replace pLux promoter - binding site: prefix + region immediately upstream the pLux promoter |
| PR_EB_166 | cagcctgcggtccggaccagtaggatcgtagaggtttacgcaagaaaatggtttgttatagtcgaataaatcgctgggacgcccg | F | Replace pLux promoter - PLux-G1: lux box with lower affinity for LuxR, promoter with same strength as the original pLux |
| PR_EB_167 | cagcctgcggtccggacctgtaggatcgtacaagtttacgcaagaaaatggtttgttatagtcgaataaatcgctgggacgcccg | F | Replace pLux promoter - Plux-76: lux box with higher orthogonality for the Las system, promoter with same strength as the original pLux |
| PR_EB_168 | cagcctgcggtccggacctgtaggatcgtacaagctgacattgtgagcggataacaatactgtttcttcaatcgctgggacgcccg | F | Replace pLux promoter - Plux54%-76: lux box with higher orthogonality for the Las system, promoter with 54% strength compared to the original pLux |
| PR_EB_175 | acctataggatcgtataggtttacgcaagaaaatggtttgttatagtcgaataaactagctaattttgtttaactttaagaaggag | F | Cloning C12 device - this primer pair amplifies mCitrine and add the pLas promoter |
| PR_EB_176 | ggaccgcaggctgaaataagacaacgttcaaatccgctcc | R | |
| PR_EB_177 | cttatttcagcctgcggtcc | F | Cloning C12 device - Amplify pTrc-LasR from G-block |
| PR_EB_183 | gaaaccacgattcagcctgc | F | Create fusion protein mCherry-LasI - this primer pair amplifies pTrc-LasI from G-block |
| PR_EB_184 | gctagcacaatacctaggactgagctagccgtcaacgggcgtcccagcgagtgtcctgtggttgagttgc | R | |
| PR_EB_206 | aggcctagcggcctgcgcagccgcgcgcttatacagctcgtccatgcc | R | Create fusion protein mCherry-LasI - binding site: the N-terminus of mCherry without the stop codon, carries HR to create a linker |
| PR_EB_207 | cgcgcggctgcgcaggccgctaggcctatgatcgttcagatcggacgc | F | Create fusion protein mCherry-LasI - binding site: the C-terminus of LasI, carries HR to create a linker |
| PR_EB_208 | ccttgcccttttttgccggattaggataccgcaaggcgctg | R | Create fusion protein mCherry-LasI - binding site: the N-terminus of LasI, it creates a stop codon |
| PR_EB_209 | tccggcaaaaaagggcaagg | F | Create fusion protein mCherry-LasI - binding site: his operon terminator, just after mCherry |
| PR_EB_313 | cgcgcggctgcgcaggccgctaggcctcttatacagctcgtccatgccg | R | Create fusion protein mCherry-LasI - this primer pair generates a different linker between mCherry and LasI |
| PR_EB_314 | aggcctagcggcctgcgcagccgcgcgatgatcgttcagatcggacgcc | F | |
| PR_EB_231 | gcgattgtactagttgcggcgaattcataggatagattctggaaacttt | F | Create a new empty backbone - this primer pair amplifies the 3 node structure from pJ1991.2 v3.2 noAraC noP(BAD), to insert it in the pCDF-SpecR |
| PR_EB_232 | gggcagggtcgttaaatagc | R | |
| PR_EB_233 | gcgattgctgaggtcctgcagagctcggacggtaaagtttcc | R | Insert C6 and C12 devices in the new backbone - this primer pair amplifies the backbone in the first node region, just after SacI restriction site |
| PR_EB_234 | ccaccgctgagcaataactaaggcttaccttactcgagcaat | F | |
| PR_EB_235 | gtgctcaatcgtgtagaaatctcttagaagaagagggtgataagcc | F | Insert C6 and C12 devices in the new backbone - this primer pair amplifies pTrc-LasR - pLas81-mCitrine fragment |
| PR_EB_236 | taagcttgtccagactattggatccgttcgcacctataggatcgtata | R | |
| PR_EB_237 | ggatccaatagtctggacaagctt | R | Insert C6 and C12 devices in the new backbone - binding site: the region upstream pLas |
| PR_EB_238 | aagagatttctacacgattgagcac | F | Insert C6 and C12 devices in the new backbone - binding site: the region downstream LasR |
| PR 277 | taattttgtttaactttaagaaggagatatacat | F | Replace pLas promoter - binding site: the RBS following pLas |
| PR_EB_252 | aagcttgtccagactattggatcccagcctgcggtccgg | F | Replace pLas promoter - Homologous to prefix and suffix, this primer pair amplifies the oligonuclotides encoding for pLas variants and carries HR for the |
| PR_EB_253 | atgtatatctccttcttaaagttaaacaaaattacgggcgtcccagcga | R | |
| PR_EB_254a | cagcctgcggtccggtgttctcgtgtgaagccattgctctgatcttttggacgtttcttcgagcctagcaagggtccgggttcaccga | F | Replace pLas promoter - this partially overlapping primer pair reconstitutes the sequence of the original pLas (iGEM registry) |
| PR_EB254b | cgggcgtcccagcgactgaagaatttatacaaatttcataattttataactagcaaatgagatagatttcggtgaacccggacccttg | R | |
| PR_EB_255 | cagcctgcggtccggacctctaggatcgtagaggtttacgcaagaaaatggtttgttatagtcgaataaatcgctgggacgcccg | F | Replace pLas promoter - pLas82 (Grant et al.) |
| PR_EB_256 | cagcctgcggtccggaactagcaaatgagatagattttacggctagctcagtcctaggtatagtgctagctcgctgggacgcccg | F | Replace pLas promoter - LasO-J23106 (Tica et al.) |
| PR_EB_257 | cagcctgcggtccggctagcaaatgagatagttacgcaagaaaatggtttgttatagtcgaataaatcgctgggacgcccg | F | Replace pLas promoter - Non cooperative consensus site (Gilbert et al.) - short |
| PR_EB_258 | cagcctgcggtccggctagcaaatgagatagtacgatttaccgaaggttctattgttacgcaagaaaatggtttgttatagtcgaataaatcgctgggacgcccg | F | Replace pLas promoter - Non cooperative consensus site (Gilbert et al.) - long (−74 to −59 from transcription initiation) |
| PR_EB_259 | cagcctgcggtccggcctgcttttctgctagttacgcaagaaaatggtttgttatagtcgaataaatcgctgggacgcccg | F | Replace pLas promoter - Cooperative consensus site (Gilbert et al.) - short |

| Name | Sequence 5' - 3' | Sense | Description |
|---|---|---|---|
| PR_EB_260 | cagcctgcggtccgg tttacggctagctcagtcctaggtatagtgctagcaactagcaaatgagatagattcgctgggacgcccg | F | Replace pLas promoter - J23106-lasO (repressible promoter, Tica et al.) |
| PR_EB_263 | cagcctgcggtccggttacgcaagaaaatggtttgttatagtcgaataaatcgctgggacgcccg | F | Replace pLas promoter - Cooperative consensus site (Gilbert et al.) - long. This is the promoter region only, the actual site is −252 to −237 from transcription initiation |
| PR_EB_264 | ctagcagaaaagcaggattcgttacaggtcagaagaattg | R | Replace pLas promoter - this primer pair amplifies the backbone to insert the Cooperative consensus site (Gilbert et al.) at positions -252 to -237 |
| PR_EB_265 | cctgcttttctgctaggtctccgattgagttttctctgc | F | |
| PR_EB_348 | cagcctgcggtccggctagcaagggtccgggttcaccgaaatctatctcatttgctagttataaaattatgaaatttgcgtaaatcttcaggatccagagaattttcgctgggacgcccg | F | Replace pLas promoter - short version of pLas81 region from Curatolo et al. |
| PR_EB_307 | cagcctgcggtccggcgtttcttcgagcctagcaagggtccgggttcaccgaaatctatctcatttgctagttataaaattatgaaatttgtataaattcttcagtcgctgggacgcccg | F | Replace pLas promoter - Prefix - pLas_iGEM_short - Suffix |
| PR_EB_308 | cagcctgcggtccggcgtttcttcgagcctagcaagggtccgggttcaccgaaatctatctcatttgctagttataaaattatgaaatttgcataaattcttcagtcgctgggacgcccg | R | Replace pLas promoter - Prefix - pLas_P.aeruginosa_short - Suffix |
| PR_EB_346 | cgggcgtcccagcgaaattggtacccctgtgtgaaattc | F | Replace pLas promoter - this primer pair amplifies the full pLas81 region from Curatolo et al., with homology regions to add prefix and suffix |
| PR_EB_347 | cagcctgcggtccggatcgatgctttacgttcacac | R | |
| PR_EB_286 | cgccttcggcgaagctagggacgagagctagcatggccctggtcgatgg | F | Include RBS and stop codons on LasR and LasI - this primer pair removes the suffix, and adds the RBS rbsKDL028-SDM |
| PR_EB_287 | gctagctctcgtccctagcttcgccgaaggcggtgtgaaattgttatccgctcac | R | |
| PR_EB_288 | tatcaccctctaagagatttctacacgattgagcac | F | Include RBS and stop codons on LasR and LasI - this primer pair removes the two extra TTC, but leaves the TAA stop codon |
| PR_EB_289 | aaatctcttagagggtgataagccccaagttg | R | |
| PR_EB_290 | cgccttcggcgaagctagggacgagagctagcatgatcgttcagatcggacgc | F | Include RBS and stop codons on LasR and LasI - Remove the suffix, and add the RBS rbsKDL028-SDM |
| PR_EB_297 | cgccttcggcgaagctagggacgagagctagcatgaaaaacatcaacgccgac | F | This primer pair replaces the RBS in between pTrc and LuxR with RBS KDL028 |
| PR_EB_298 | gctagctctcgtccctagcttcgccgaaggcggtgtgaaattgttatccgctcac | R | |
| PR_EB_319 | ttggaaactgaagatgcggc | F | Clone the C6-C12 device - binding site: before the terminator of the 3° node in the 3-node backbone |
| PR_EB_320 | aacgacgagtgtaataggacg | R | Clone the C6-C12 device - this primer pair amplified the 3-node backbone structure in the region of the 3° node |
| PR_EB_321 | cgccttcggcgaagc | F | |
| PR_EB_322 | gccgcatcttcagtttccaattaggataccgcaaggcgctg | R | Clone the C6-C12 device - binding site: the N-terminus of LasI, with homology region for the 3° node region in the 3-node backbone (EB_319) |
| PR_EB_323 | cgtcctattacactcgtcgttcagcctgcggtccgg | F | Clone the C6-C12 device - this primer pair amplifies pLux76 with homology regions for the 3-node backbone and for LasI (EB_320 and EB_321) |
| PR_EB_324 | ccctagcttcgccgaaggcgcgggcgtcccagcga | R | |
| PR_EB_335 | ggcatggacgagctgtataag | F | Clone the C6-C12-device - binding site: the N-terminus of mCherry, amplify all the his operon terminator |
| PR_EB_338 | gaaaagaatatgctcgttcccg | F | Clone the C6-C12-device - binding site: between the mCherry terminator and the pTrc promoter, to amplify pLux-mCherry and pTrc-LuxR, respectively |
| PR_EB_339 | cgggaacgagcatattcttttc | R | |
| PR_EB_363 | gcagggtcgttaaatagccgaactgggtgggactcggc | R | Check for cross talk between C6 and C12 - Remove the LasR-mCitrine part. |
| PR_EB_364 | gccgcatcttcagtttccaaaactgggtgggactcggc | R | |
| PR_EB_365 | taattttgtttaactttaagaaggagatatacatatggtgagcaagggcgagg | F | Check for cross talk between C6 and C12 - this primer pair replaces the pLux76 in between LasI and mCherry with an RBS |
| PR_EB_366 | atgtatatctccttcttaaagttaaacaaaattaggaagtatcgtagttcacccaatcag | R | |
| PR_EB_371 | cgccttcggcgaagctagggacgagagctagcatgattgagaatacctatagcgaaaagttcg | F | Replace Las components with Cin components - this primer pair amplifies cinR with homology arms to replace LasR in the backbone of C6_device |
| PR_EB_372 | gtgctcaatcgtgtagaaatctcttctattaccaattacgtcgcgtcatgc | R | |
| PR_EB_373 | aagcttgtccagactattggatccttttgtttctggtctcccgc | F | Replace Las components with Cin components - this primer pair amplifies pCin with homology arms to replace pLas81 in the backbone of C6_device |
| PR_EB_374 | ctagtatttcccctctttctctagtattaaac | R | |
| PR_EB_375 | gtttaatactagagaaagaggggaaatactagatgtctaaaggtgaagaattattcactgg | F | Replace Las components with Cin components - binding site: the C-terminus of mCitrine, with homology region for the pCin-EYFP RBS |
| PR_EB_376 | cggcgcctccctatcagt | R | Replace Las components with Cin components - binding site: RBS-KDL027 |
| PR_EB_377 | atggtgagcaagggcgagg | F | Replace Las components with Cin components - binding site: the C-terminus of mCherry |
| PR_EB_378 | atgtatatctccttcttaaagttaaacaaaattatttattacgctgcaagggcg | R | Replace Las components with Cin components - this primer pair amplifies cinI with homology regions to be inserted between pLux76 and mCherry |
| PR_EB_379 | actgatagggaggcgccgatgcgggggagtgttactag | F | |
| PR_EB_380 | aagagatttctacacgattgagcaccggatttgaacgttgtgaagc | F | Replace Las components with Cin components - Opens C6_device to insert cinR just before the rrnB T2 terminator |
| PR_EB_396 | ccacggcatggacgaactgtataaataataacagtagtacttgcacagacgtc | R | Remove LuxI from GFP-LuxI fusion protein - Add 2x TAA stop codons at the N-terminus of GFP |
| PR_409 | tttatacagttcgtccatgccgtgg | F | Remove LuxI from GFP-LuxI fusion protein - binding site: the N-terminus of sfGFP |
| PR_EB_397 | cagcctgcggtccggacctgtaggatcgtacaggtttacttgtgagcggataacaatatagtgtgtggaattgtgagcggataacaatttcgctgggacgcccg | F | Replace pLux76 with pLuxLac - pLuxLac sequence |
| PR_EB_399 | tgcaggacctcagcaatcgcttggacagcctgcggtccgg | F | Replace pLux76 with pLuxLac - this primer pair amplifies PR_EB_397 with homology regions |
| PR_EB_400 | cagttgaacgatagttaatactcgggcgtcccagcga | R | |

TABLE VII. Equation and parameters of constrained logistic curves fitted to experimental points in Figure 2c. Equation: $y = \frac{100}{1+e^{-k\cdot(x-x_0)}}$. Method: Levenberg-Marquardt algorithm.

| **Set of data** | Initial state: green | | Initial state: mixed | | Initial state: blue | |
|---|---|---|---|---|---|---|
| | $k$ | $x_0$ | $k$ | $x_0$ | $k$ | $x_0$ |
| Green percentage | 1.171 | -2.251 | 0.514 | -0.594 | 0.197 | 13.996 |
| Blue percentage | -1.207 | -2.379 | -0.541 | -0.756 | -0.200 | 13.303 |

TABLE VIII. Equation and parameters of exponential curves fitted to experimental points in Figure 4b. Equation: $y = y_1 + (y_0 - y_1) \cdot e^{-k\cdot x}$. Method: Trust Region Reflective algorithm.

| **Set of data** | $k$ | $y_0$ | $y_1$ |
|---|---|---|---|
| Reporter strain (white hexagons) | 0.20 | 390.81 | 306.34 |
| Reporter strain + C6-HSL (red circles) | 0.20 | 6054.56 | 303.30 |
| Reporter strain + sender cells (orange triangles) | 0.50 | 1221.08 | 190.39 |

TABLE IX. Equation and parameters of exponential curves fitted to experimental and simulation points in Figure 5b,c. Equation: $y = y_1 + (y_0 - y_1) \cdot e^{-k\cdot x}$. Method: Trust Region Reflective algorithm.

| **Set of data** | $k$ | $y_0$ | $y_1$ |
|---|---|---|---|
| Experimental points (Figure 5b) | 0.995 | 1231.224 | 232.860 |
| Simulation points (Figure 5c) | 1.082 | 0.090 | 0.0001 |

TABLE X. Equation and parameters of exponential curves fitted to experimental points in Figure 7e. Equation: $y = y_1 + (y_0 - y_1) \cdot e^{-k\cdot x}$. Method: Levenberg-Marquardt algorithm.

| **Set of data** | $k$ | $y_0$ | $y_1$ |
|---|---|---|---|
| Red fluorescence intensity | 1.781 | 18906.42 | 2614.911 |
| Yellow fluorescence intensity | 1.650 | 10107.657 | 1132.947 |

TABLE XI. Equation and parameters of stretched exponential curves fitted to simulation points in Figure S9. Equation: $y = y_1 + (y_0 - y_1) \cdot e^{(-k \cdot x)^{\beta}}$. Method: Trust Region Reflective algorithm.

| **Set of data** | $k$ | $\beta$ | $y_0$ | $y_1$ |
|---|---|---|---|---|
| Reporter strain (black hexagons) | 0.5 | 1.2 | 0.0 | 0.0 |
| Reporter strain + C6-HSL (red circles) | 0.138 | 2.213 | 34742.707 | 0.01 |
| Reporter strain + sender cells (orange triangles) | 1.0 | 1.013 | 1240.274 | 0.01 |

TABLE XII. Equation and parameters of power-law curves fitted to simulation points in Figure S15b. Equation: $y = \frac{A}{(x+x_0)^p}$. Method: manual smoothing.

| **Set of data** | $A$ | $x_0$ | $p$ |
|---|---|---|---|
| Leftmost panel - mCherry | $5 \cdot 10^9$ | 2 | 13 |
| Leftmost panel - mCitrine | $1.4 \cdot 10^{11}$ | 4.8 | 10 |
| Central panel - mCherry | $6 \cdot 10^{10}$ | 4.5 | 11 |
| Central panel - mCitrine | $4 \cdot 10^{10}$ | 6 | 10 |
| Rightmost panel - mCherry | $4 \cdot 10^{10}$ | 6 | 10 |
| Rightmost panel - mCitrine | $2 \cdot 10^{14}$ | 11 | 12 |
