## Supplementary material for "An engineered multi-step differentiation program in *Escherichia coli* for self-organized spatial patterning": Description supplementary movies

**Supplementary Movies 1, 2, 3, 4: microscopy time-lapses of the 2-step sequential differentiation system in different conditions.** Cells were pre-cultured starting from the blue state with 1 mM IPTG (Movies 1, 2, 3) and starting from the green state with 9 $\mu$M IPTG (Movie 4), then plated on solid surface in absence of chemical inducers. The plate was then inoculated in the centre with 2 $\mu$l of 3O-C6-HSL 50 $\mu$M immediately before incubation (Movie 1), with 2 $\mu$l of 3O-C6-HSL 50 $\mu$M 8 hours after incubation started (Movie 2) or with 1 $\mu$l of green sender cells (pre-differentiated with 100 nM aTc) at OD=1 immediately before incubation (Movie 3). In the case of the intermixed population, no additional positional information was added (Movie 4).

**Supplementary Movie 5: a spatio-temporal simulation of the spatial pattern generated by the 2-step sequential differentiation system.** Colonies are colour-coded, from left to right, according to: TS state (green-senders, blue-receivers), mCherry fluorescence intensity, full pattern.

**Supplementary Movie 6: diffusion wavefronts of 3O-C6-HSL and 3O-C14-HSL in simulations of the 3-step sequential differentiation system.** A single green-sender colony was simulated in the centre of the plate, surrounded by blue receivers. The green and red lines represent the radial profiles of 3O-C6-HSL and 3O-C14-HSL respectively, from the centre to the periphery of the plate (x axis, 0 corresponds to the position of the green-sender colony), over time (151 simulation frames distributed from 9 h to 24 h). The wavefront of 3O-C14-HSL is always behind the wavefront of 3O-C6-HSL.
