## Supplementary figures and images for "An engineered multi-step differentiation program in *Escherichia coli* for self-organized spatial patterning"

### Supplementary movie 5

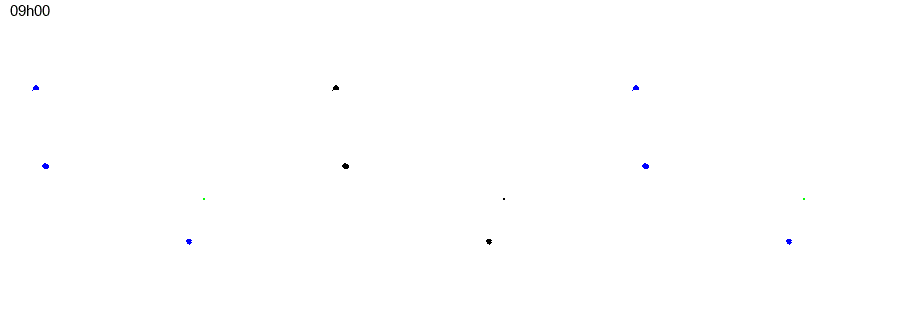

### Supplementary movie 6

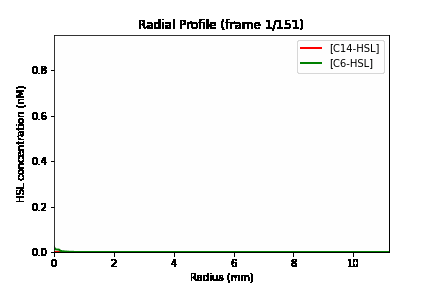
